# Drug-centric knowledge graph: representing therapeutic modality, pharmacology and multiscale biology

**DOI:** 10.64898/2026.10.15.759051

**Authors:** Alon Bar, Michael Shapiro, Gilad Erez, Doron Levinson, Tal Raisfeld, Uri Shamgar, Amichai Perlman, Omri Matalon, Flavio Dormont

**Affiliations:** QuantHealth Ltd., Tel Aviv, Israel

**Keywords:** biomedical knowledge graph, drug-centric knowledge graph, therapeutic modality, compound identity, antibody–drug conjugates, pharmacokinetics, graph representation learning

## Abstract

Biomedical knowledge graphs link targets to mechanisms, diseases and phenotypes, but target-centered representations omit much of what distinguishes therapeutic interventions. We introduce a drug-centric knowledge graph, whose identity model, schema and information-selection priorities are organized around the therapeutic intervention. Each intervention has one identity, linked through relation types shared by all compounds, including target actions, indications, adverse effects, classifications, half-life and direct links to pathways, Gene Ontology concepts, cells and anatomy, and through modality-specific relation types describing small-molecule chemistry, antibody sequences and formats, protein-therapy sequences and antibody–drug conjugate components. Integrating 40 source resources, the graph contains 166,261 nodes of 27 types, 8,106,906 directed edges of 129 relation types and 18,429 compounds in eight therapeutic categories. Construction combines source-specific extraction, compound-identity resolution, language-model-assisted vocabulary and predicate mapping, confidence scoring and connectivity filtering; an expert-reviewed workflow adds investigational therapeutics without altering the base graph. The design offers a reusable template for representing interventions as structured, multimodal objects.

## Background & Summary

Drug development requires the integration of evidence across molecular, physiological and clinical scales. A therapeutic intervention has a physical composition, interacts with biological systems and produces effects that depend on its exposure and the context of administration. Information describing these dimensions is distributed across compound databases, regulatory documents, molecular interaction resources, disease ontologies and the scientific literature. Biomedical knowledge graphs integrate these heterogeneous sources through entities and typed relationships, supporting mechanistic investigation, information retrieval and machine learning.

Existing graphs emphasize complementary aspects of this integration (Table 1). Hetionet connects compounds, diseases and molecular biology to support drug-repurposing hypotheses, whereas PrimeKG organizes multimodal knowledge for precision medicine and includes clinical drug features [1, 2]. The Biological Insights Knowledge Graph describes an organizational resource spanning data integration and multiple drug-development applications [3], and OptimusKG combines a schema-constrained property graph with type-specific metadata, provenance and an explicit evidence-support assessment [4]. Drug-focused resources provide further precedents: PharmKG combines relational structure with chemical, gene-expression and disease features for representation learning [5]; OREGANO integrates drug and natural-compound information for computational repurposing [6]; DrugMechDB curates mechanistic paths between interventions and indications [7]; and NP-KG represents pharmacokinetic mechanisms underlying natural product–drug interactions [8].

**Table 1.** Drug-centric knowledge graphs in the landscape of biomedical knowledge graphs. The comparison summarizes representative scientific emphases; resources may support capabilities beyond those listed.

| Resource | Published representational emphasis | Relevance to the drug-centric design |
| --- | --- | --- |
| Hetionet [1] | Integrated compounds, diseases and biological entities for drug-repurposing hypotheses | Establishes the shared biomedical neighborhood as a basis for therapeutic reasoning |
| PrimeKG [2] | Disease-oriented multimodal integration with clinical drug features, including half-life | Drug attributes and pharmacokinetic information complement relational topology |
| BIKG [3] | Organizational integration, graph lifecycle and multiple drug-development applications | Motivates documenting construction, maintenance and downstream consumption together |
| OptimusKG [4] | Schema-constrained property graph, type-specific metadata, provenance and literature-support assessment | Makes type-specific information, provenance and evidence support inspectable beyond graph connectivity |
| PharmKG [5] | Chemical, gene-expression and disease features integrated with relational learning | Feature-aware learning requires explicit treatment of information beyond entity identifiers |
| OREGANO [6] | Drug and natural-compound integration for computational repurposing | Establishes rich drug characterization as a prior resource-design objective |
| DrugMechDB [7] | Curated paths describing drug mechanisms leading to indications | Direct mechanistic context is an established complement to target lists |
| NP-KG [8] | Pharmacokinetic natural product–drug interaction knowledge | Pharmacokinetic graph representation has task-specific semantics |
| This work | Shared therapeutic identity, modality-specific components, half-life and qualitative pharmacology, linked to multiscale biology | Combines shared therapeutic identity with modality-specific composition, pharmacology and biological context |

These resources establish the value of molecular features, pharmacological context and mechanistic relationships. The representational challenge addressed here is to preserve the features that distinguish interventions while retaining a common basis for comparison across therapeutic classes. Two antibodies may recognize the same target yet differ in sequence, format, isotype or half-life. Two antibody–drug conjugates (ADCs) may share an antibody but differ in linker or payload. Conversely, therapeutics with different molecular architectures may converge on related pathways or clinical indications. A target-only description compresses these distinctions, while isolated modality-specific datasets fragment the biological context needed to compare them.

We use the term **drug-centric knowledge graph** for a biomedical graph whose identity model, schema and information-selection priorities are organized around the therapeutic intervention. Three design principles follow. First, the complete intervention is the unit of identity: each therapeutic receives a single identifier, and its chemical structures, sequences and components are linked to that identity rather than defining it; when a component is itself a therapeutic, such as the antibody of an ADC, the composite links to the component’s own identity. Second, the schema has two tiers: relation types shared by all compounds, such as targets, indications, adverse effects, classifications and biological context, provide a common basis for comparison, whereas modality-specific relation types describe the composition of each therapeutic class. Third, relation semantics separate molecular recognition, functional action and drug disposition, so that comparisons can distinguish binding from functional effect and from metabolism or transport. Together, these principles allow a graph to represent what a drug is, how it acts and where it acts. Therapeutic modality refers to the intervention class, such as a small molecule or antibody; data modality refers to the form of the information, such as a chemical structure, sequence, categorical annotation or numerical value. The design does not depend on a particular construction framework and complements adaptable frameworks such as BioCypher and feature-aware graph-learning approaches [5, 9].

Here we describe a graph that implements this design. It organizes information into three connected domains, namely compound characteristics, the biological landscape and clinical context, around a common Compound identity (Fig. 1). Half-life annotations provide pharmacokinetic (PK) information, while binding and functional-action relationships contribute pharmacodynamic (PD) context, connecting the resource to the distinction between drug exposure and biological response [10, 11]. We report the data model, the construction pipeline and its harmonization methods, the composition of the resulting graph and the quality controls applied during its assembly.

**Figure 1.**
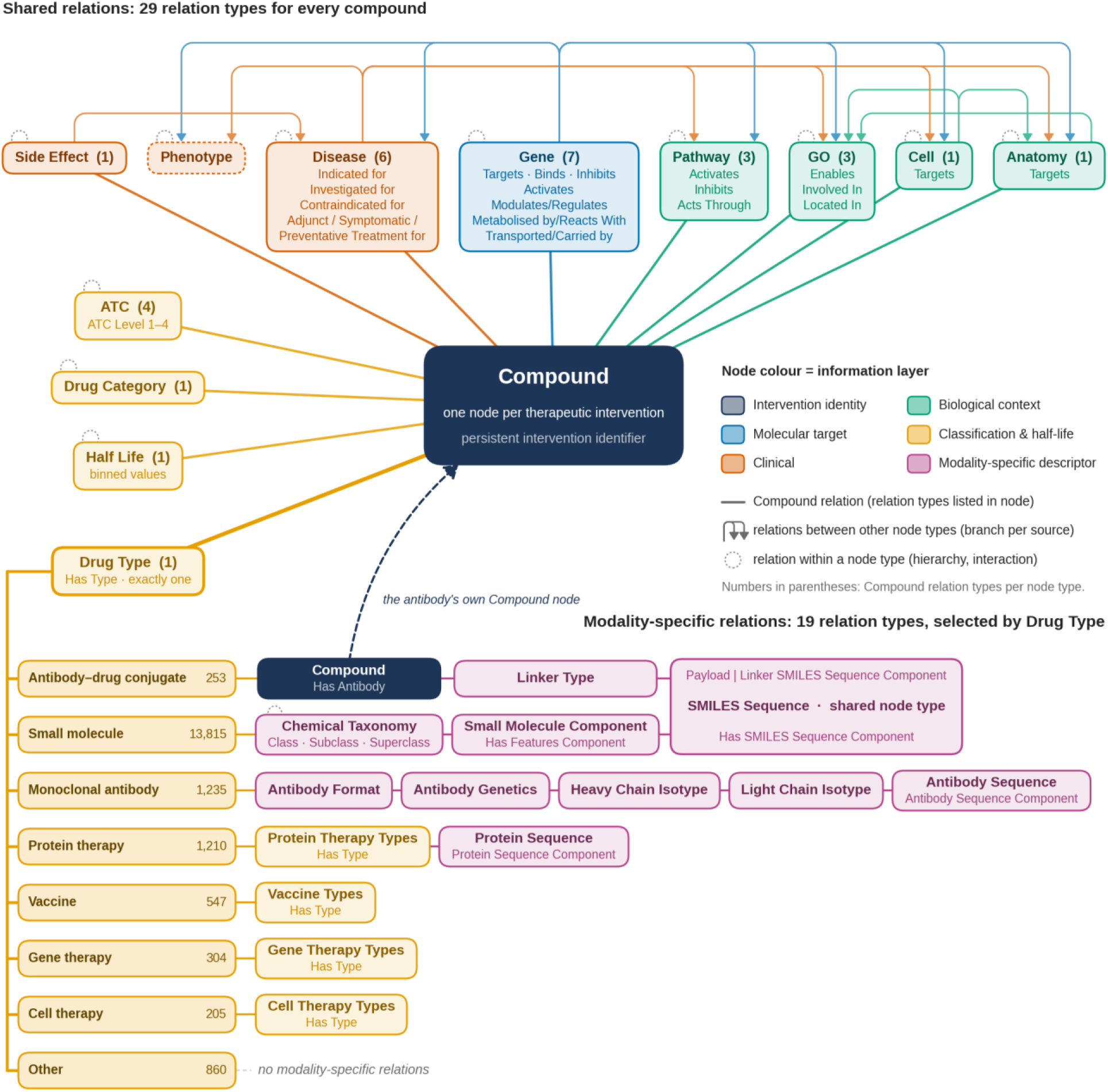
Drug-centric data model. Each therapeutic intervention is represented by one Compound node with a persistent intervention identifier. Lines from the Compound node show the 29 relation types shared by all compounds; numbers in parentheses give the relation types per target node type, and relation names are listed in each node. Branches above the top row show relations between other node types, coloured by source node type with arrowheads marking direction, and dotted loops mark relations within a node type, such as hierarchies and interactions; Phenotype has no direct compound relation. Bottom, exactly one Has Type relation assigns each compound to one of eight therapeutic categories (compounds per category shown at the right of each label), which determines the applicable modality-specific relation types (19 in total). Each descriptor node type attaches to the compound separately; relations named Has <node type> are shown by the node type alone. SMILES Sequence is one node type used by both small molecules and ADCs, and the Has Antibody relation of an ADC points to the Compound node of its antibody. Colours indicate information layers.

## Methods

### Drug-centric data model

Each therapeutic intervention is represented by a single Compound node carrying a persistent intervention identifier. The data model attached to this identifier has three parts (Fig. 1; Table 2): relation types shared by all compounds; exactly one Drug Type, which assigns the intervention to one of eight therapeutic categories; and modality-specific relation types determined by that category. A compound can have several edges of the same relation type, for example to multiple molecular targets or indications. The 29 shared relation types connect compounds to genes, diseases, side effects, pathways, Gene Ontology (GO) concepts, cells, anatomical structures, drug categories, Anatomical Therapeutic Chemical (ATC) classes, Drug Types and half-life bins. The 19 modality-specific relation types describe small molecules by chemical taxonomy, simplified molecular-input line-entry system (SMILES) sequence components and feature components; monoclonal antibodies by sequence components, format, genetics and heavy- and light-chain isotypes; protein therapies by sequence components and therapy type; ADCs by their antibody, linker and payload; and vaccines, gene therapies and cell therapies by a type classification.

**Table 2.** Compound relation schema. Shared relation types apply to every compound; modality-specific relation types apply according to the compound’s Drug Type. The schema defines 29 shared and 19 modality-specific relation types, and a compound can have several edges of the same type. Relation identifiers in this article take the form Source::Target::Predicate, for example Compound::Gene::Binds; the edge table serializes the same relation as Binds::Compound:Gene. Sources are described in Table 3, and the full schema is provided in Supplementary Data 1. SMILES, simplified molecular-input line-entry system; ATC, Anatomical Therapeutic Chemical.

| Target node type | Relation types (predicates) | Principal sources |
| --- | --- | --- |
| **Shared Compound relations (all therapeutic categories)** |  |  |
| Gene | Targets; Binds; Inhibits; Activates; Modulates/Regulates; Metabolised by/Reacts With; Transported/Carried by | ADCdb; DGIdb; DrugBank; IntAct; OpenFDA; Open Targets; SAbDab; SIGNOR; TTD |
| Disease | Indicated for; Investigated for; Contraindicated for; Adjunct Treatment for; Symptomatic Treatment for; Preventative Treatment for | ADCdb; DrugBank; EMA; OpenFDA; SAbDab; TTD |
| Side Effect | Has Side Effect | DrugBank; OnSIDES; OpenFDA |
| Drug Type | Has Type (exactly one per compound) | DrugBank; ADCdb |
| Drug Category | Has Drug Category | DrugBank; MeSH |
| ATC | ATC Level 1; ATC Level 2; ATC Level 3; ATC Level 4 | DrugBank |
| Half Life | Has Half Life | DrugBank (binned values) |
| Pathway | Activates; Inhibits; Acts Through | Reactome; Wikipedia; PubMed |
| GO | Enables; Involved In; Located In | Wikipedia; PubMed |
| Anatomy | Targets | Wikipedia; PubMed |
| Cell | Targets | Wikipedia; PubMed |
| **Small molecule** |  |  |
| Chemical Taxonomy | Class; Subclass; Superclass | DrugBank |
| SMILES Sequence | Has SMILES Sequence Component | DrugBank |
| Small Molecule Component | Has Features Component | DrugBank |
| **Monoclonal antibody** |  |  |
| Antibody Sequence | Antibody Sequence Component | SAbDab |
| Antibody Format; Antibody Genetics | Has Antibody Format; Has Antibody Genetics | SAbDab; ADCdb |
| Heavy Chain Isotype; Light Chain Isotype | Has Heavy Chain Isotype; Has Light Chain Isotype | SAbDab; ADCdb |
| **Protein therapy** |  |  |
| Protein Sequence | Protein Sequence Component | DrugBank; internal curated tables |
| Protein Therapy Types | Has Type | DrugBank |
| **Antibody–drug conjugate** |  |  |
| Compound (monoclonal antibody) | Has Antibody | ADCdb |
| SMILES Sequence | Payload SMILES Sequence Component; Linker SMILES Sequence Component | ADCdb |
| Linker Type | Has Linker Type | ADCdb |
| *Vaccine, gene therapy and cell therapy* |  |  |
| Vaccine Types; Gene Therapy Types; Cell Therapy Types | Has Type | DrugBank |

This organization separates the complete intervention from the information used to describe it. Modality-specific descriptors are represented as typed nodes, so that a sequence, chemical component or linker type can contribute to a drug representation and connect the interventions that share it without defining an additional therapeutic intervention. For composite therapeutics, the distinction also separates component similarity from whole-drug identity: an ADC links through Has Antibody to the Compound node of its monoclonal antibody, which carries its own antibody descriptors, and through component relations to its linker and payload structures. Two ADCs can therefore be compared at the antibody, linker or payload level while retaining distinct identities and clinical associations.

### Construction workflow

The graph was assembled with custom, source-specific Python notebooks and package functions executed on Databricks; no off-the-shelf graph-construction framework was used. Construction proceeds in six stages (Fig. 2): (1) download and preprocessing of raw source files; (2) generation of standard vocabularies, intervention identifiers, entity mappers and component models; (3) extraction of source triplets under extraction contracts, which specify for each source the required columns, the permitted source and target node types and the approved identifier vocabularies; (4) transformation of triplets to standard identifiers and registered predicates; (5) validation, comprising structural checks and confidence scoring; and (6) post-processing and graph assembly. Extract snapshots are stored as Parquet files, intermediate graph data as Delta tables and per-drug records as JSON documents. Stages 5 and 6 are described under Technical Validation. A separate enrichment workflow produces reviewed information about investigational therapeutics without altering the base graph.

**Figure 2.**
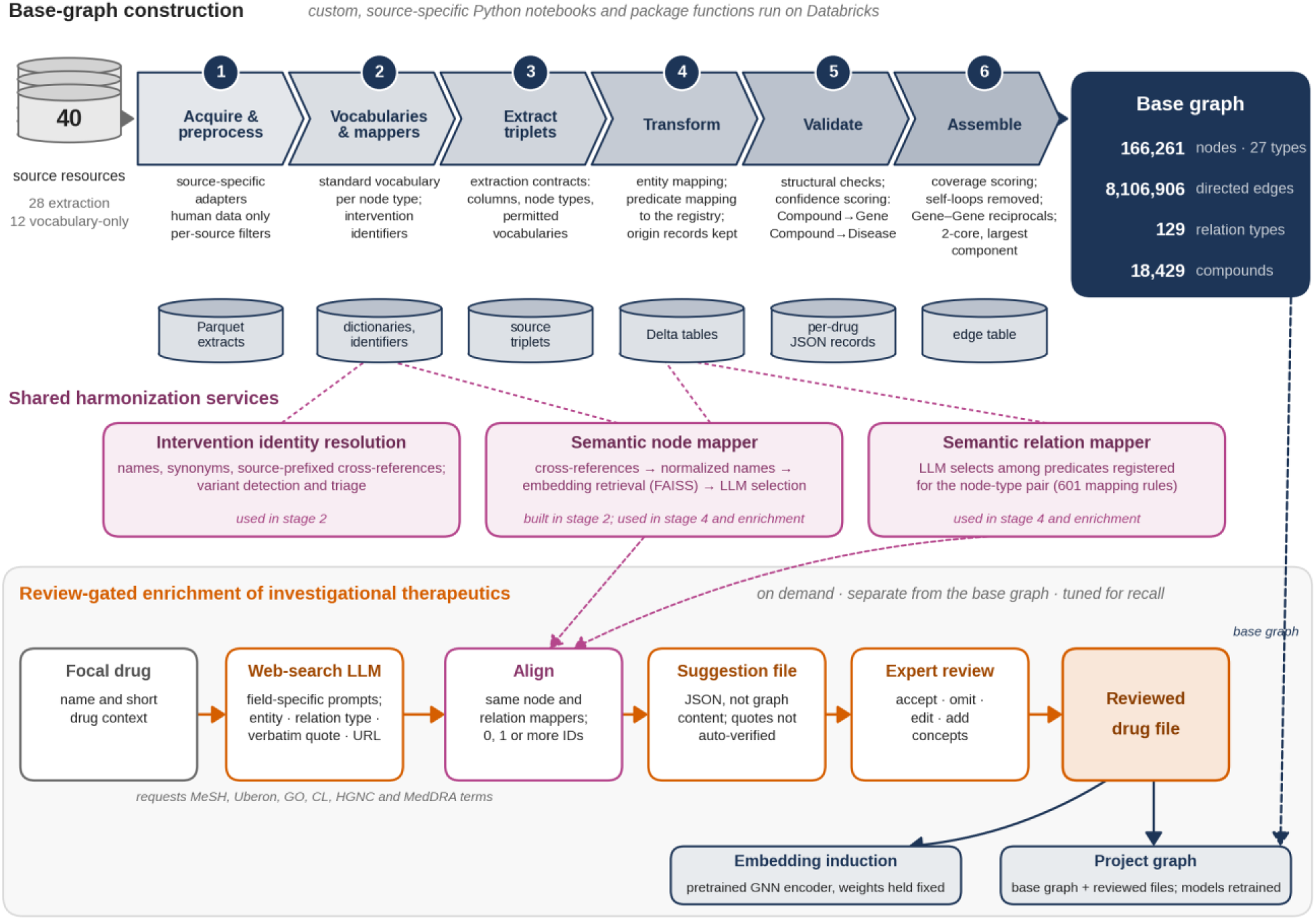
Construction of the graph. Top, the six stages of base-graph construction with the principal operations of each stage and the data artefacts it produces, ending in the post-processed base graph. Middle, harmonization services shared by construction and enrichment. Bottom, the on-demand enrichment workflow for investigational therapeutics: web-search-enabled language models propose relations with verbatim supporting quotations and source URLs, proposals are aligned with the same node and relation mappers, and expert review produces a reviewed drug file kept separate from the base graph. Reviewed files are used either to induce an embedding with a pretrained graph neural network (GNN) encoder whose weights are held fixed or to build a project graph on which models are retrained.

### Source resources and extraction

The graph draws on 40 source resources: curated biomedical databases, ontologies and controlled vocabularies, regulatory resources and text from Wikipedia and PubMed (Table 3; Fig. 3). Twelve resources are used only for vocabulary construction and identifier mapping, and 28 are designated extraction sources, of which 26 contribute relations to the graph; the other two, DOID and PubChem, supply alternate disease identifiers and compound cross-references. Codes from vocabularies that were not acquired separately, namely OMIM and SNOMED, which are obtained through UMLS, and ATC, which is derived from DrugBank, are not listed as separate sources. A single resource can contribute to several domains: GO, for example, supplies biological-process, molecular-function and cellular-component concepts, while regulatory sources contribute both indication and safety information.

**Figure 3.**
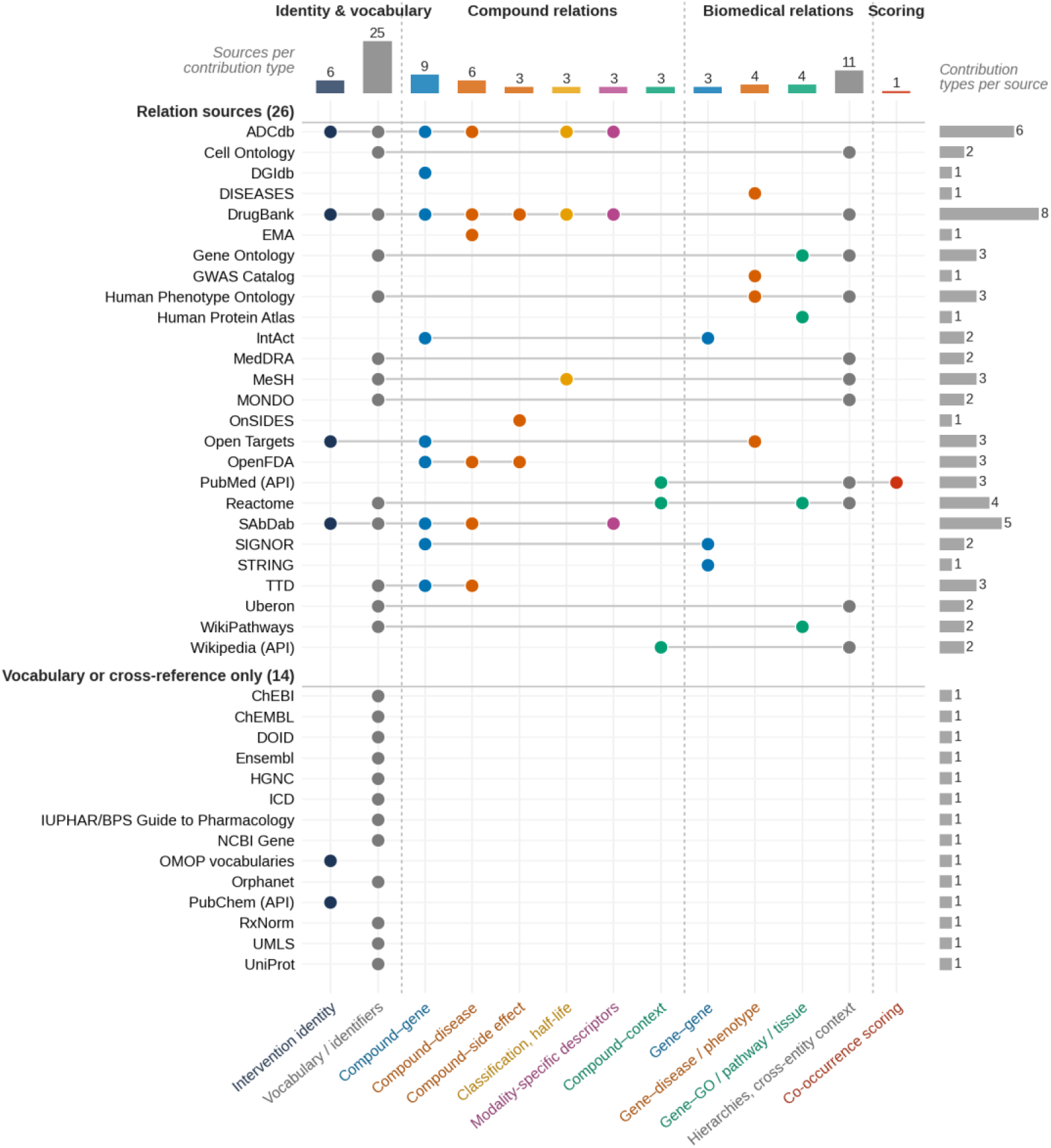
Contributions of the source resources. Each row is one of the 40 resources in Table 3 and each column a type of contribution; lines connect the contributions of one resource. Resources are grouped into those contributing relations to the graph (26) and those used only for vocabularies, identifiers or cross-references (14). Bars give the number of resources per contribution type (top) and the number of contribution types per resource (right). API, accessed through an application programming interface.

**Table 3.**
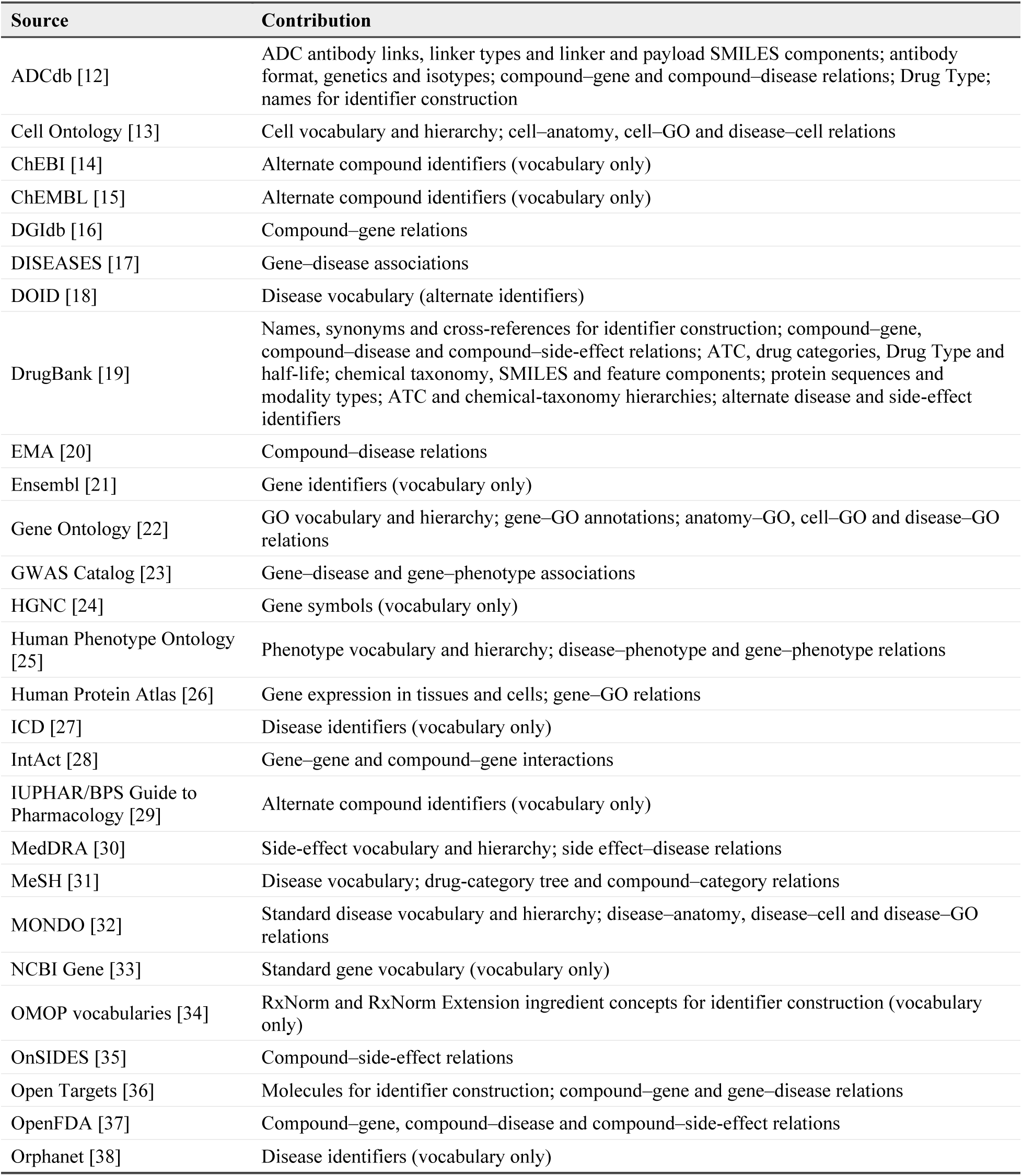
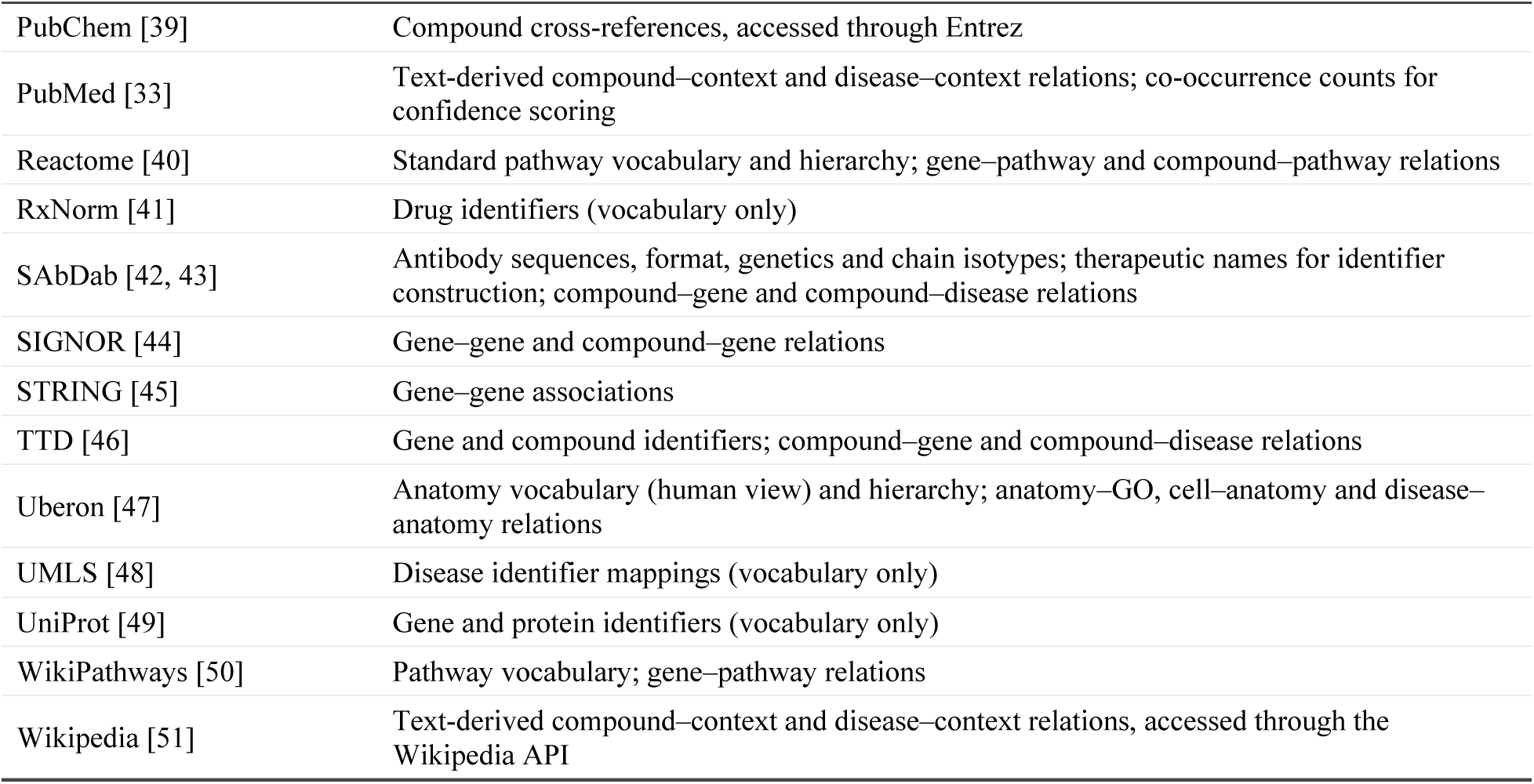
Source resources. Contribution summarizes the vocabularies and relations to which each resource contributes. PubChem, PubMed and Wikipedia were accessed through application programming interfaces. Antibody and linker content from SAbDab and ADCdb was integrated through internally curated tables, which also contribute protein-therapy sequences. The complete inventory is provided in Supplementary Data 1.

Each source is processed by a custom, source-specific Python adapter, and extraction is restricted to human data. Inclusion filters are defined per source rather than by a common quality policy: for example, Open Targets gene–disease associations are retained when their association score exceeds 0.2 [36]; Human Protein Atlas expression records are retained when their confidence is medium or high and their score exceeds 0.05 [26]; and DISEASES associations are taken from the human disease-association file with one contributing source excluded [17]. Direct compound–GO, compound–cell and compound–anatomy relations, compound–pathway relations not provided by Reactome and several disease-context relations are derived by text extraction from Wikipedia articles and PubMed records. No confidence cutoffs are applied to text-derived relations; like other relations outside the scored compound–gene and compound–disease subset, they are subject to the structural checks and assembly filters described under Technical Validation.

Source integration must also preserve the meaning of the evidence. Regulatory indications, curated disease associations and literature-derived mechanistic statements describe different relationships between a drug and its clinical or biological context. Similarly, agreement between databases can arise from reuse of a common underlying observation. These distinctions motivate source-aware integration and the interpretation of graph relationships in their original evidential context.

### Text extraction of compound–context relations

Compound–pathway, compound–GO, compound–anatomy and compound–cell relations are extracted from text by a large language model (OpenAI GPT-4.1, temperature 0) that returns structured output. For each compound, the extractor assembles two kinds of context: the English Wikipedia article for the compound [51], when one is found, with its title, URL, summary and full text; and up to ten PubMed abstracts [33] retrieved by relevance-ranked search without a publication-date restriction, supplied in batches of five with their identifier, title, journal and year. Each context is processed with three task instructions, covering pathway relations (activates, inhibits, acts through), GO relations (enables, involved in, located in) and anatomical and cellular targets, under a shared instruction to extract only statements made explicitly about the named compound, to prefer Gene Ontology, Uberon, Cell Ontology and Reactome terminology, and to return a supporting text reference. The task instructions restrict pathway and GO terms to the sentence that names the compound or its immediate context, exclude annotation lists and background statements, exclude sites of metabolism or excretion from anatomical targets, and ask the model to check that GO terms and targets are consistent with the compound’s known therapeutic class or indication (Supplementary Data 2). The outputs of all contexts are merged, and their entities and relation labels are aligned with the node and relation mappers described below. These are the retrieval settings of the construction code rather than a record of the articles processed for the reported build.

### Compound identity resolution

Intervention identifiers are constructed from DrugBank names, synonyms and external identifiers [19]; OMOP RxNorm and RxNorm Extension ingredient concepts [34, 41]; therapeutic names from SAbDab [42, 43] and ADCdb [12]; and Open Targets molecules [36] (Fig. 4a). Candidate records are matched by comparing normalized names and synonyms together with source-prefixed cross-references, including DrugBank and PubChem Compound identifiers [39]; PubChem Substance identifiers are excluded from this route. Identifiers from the IUPHAR/BPS Guide to Pharmacology [29], ChEBI [14] and ChEMBL [15] are retained as alternate compound identifiers. Overlapping candidates can then be merged. The procedure also detects some salt, solvate, strength, formulation, biosimilar and development-code variants and triages mixed or out-of-scope entries. Identity assignments were checked by manual, qualitative review. These rules are implementation heuristics rather than a validated identity policy covering every salt, stereoisomer, active moiety, brand, mixture, formulation or biologic construct, and the error rate of identity resolution has not been quantified.

**Figure 4.**
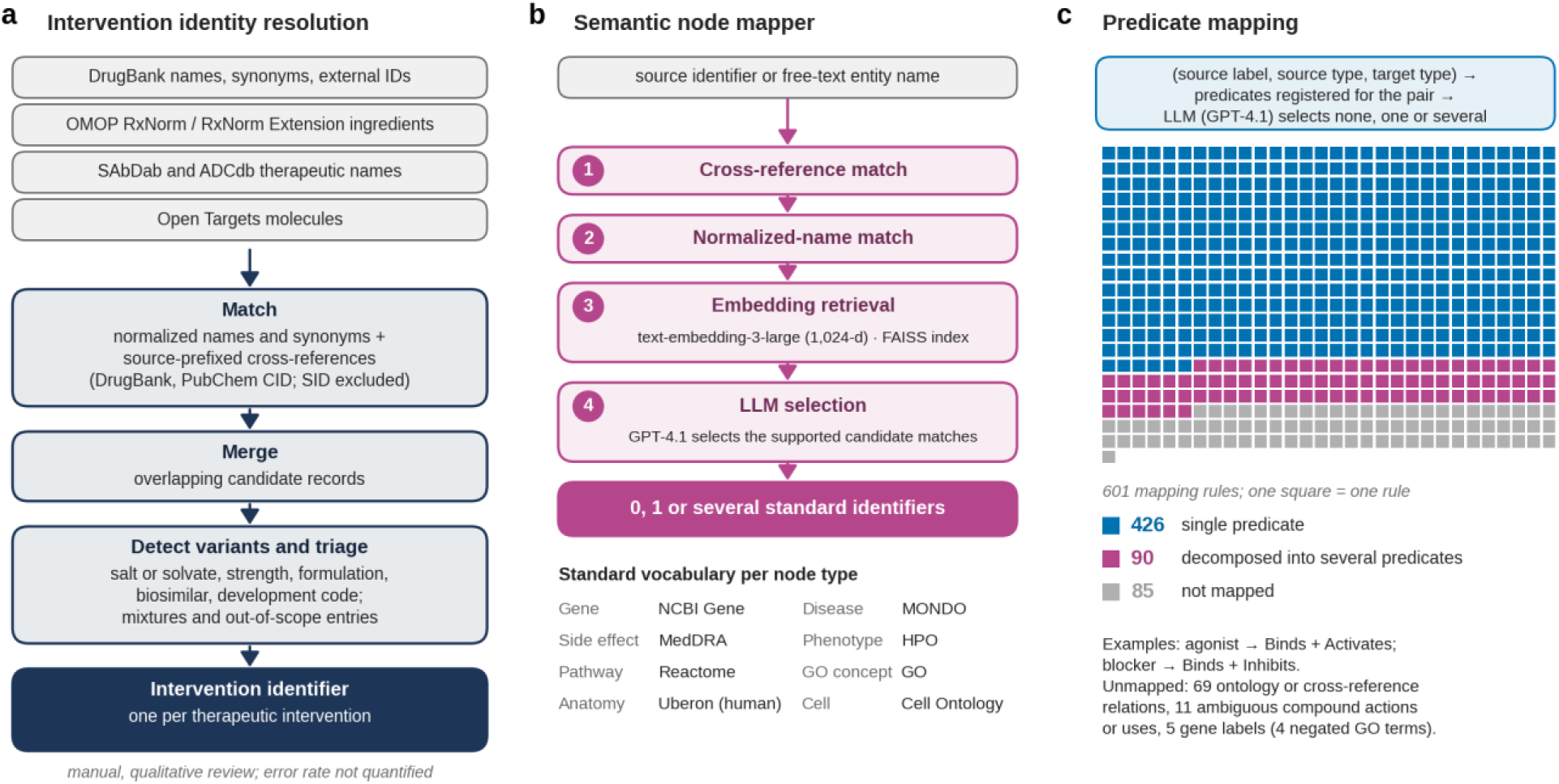
Harmonization of identities, entities and predicates. **a**, Construction of intervention identifiers from five input sources by matching, merging, variant detection and triage. **b**, The semantic node mapper resolves a source identifier or free-text name by cross-reference and normalized-name matching and, for unresolved text, by embedding retrieval followed by language-model selection; a term can map to zero, one or several standard identifiers. The standard vocabulary of each biomedical node type is listed below. **c**, Predicate mapping. The relation mapper selects among the predicates registered for a source–target node-type pair; the 601 rules of the predicate-mapping table (one square per rule) map a source relation label to a single predicate (426), to several predicates (90) or to none (85).

The central objective of identity resolution is to consolidate equivalent descriptions while preserving distinctions that affect therapeutic interpretation. Salts, formulations, mixtures and biologic constructs can represent different levels of identity; treating all such records as interchangeable can obscure clinically relevant differences, whereas retaining equivalent records as separate interventions duplicates evidence. Compound identity therefore defines both the unit of integration and the unit of analysis in downstream tasks.

### Vocabulary normalization and entity mapping

Each biomedical node type has a standard vocabulary: NCBI Gene for genes, MONDO for diseases, MedDRA for side effects, the Human Phenotype Ontology for phenotypes, Reactome for pathways, GO for GO concepts, the human view of Uberon for anatomy and the Cell Ontology for cells; other resources, such as Orphanet [38] for diseases, supply alternate identifiers, names and synonyms. Drug Types, half-life bins and the modality-type taxonomies are defined internally. A semantic node mapper links source identifiers and free-text entity names to these vocabularies (Fig. 4b). It first checks cross-references and normalized names; for unresolved text, it retrieves candidate vocabulary entries from a FAISS index [52] of OpenAI text-embedding-3-large embeddings (1,024 dimensions), and a large language model (OpenAI GPT-4.1) selects the supported matches. A term can map to zero, one or several standard identifiers. Vocabulary alignment connects heterogeneous source labels to shared concepts, whereas typed relations specify how a drug is associated with them: an anatomical site of action, expression of a molecular target in a tissue and measured drug distribution are distinct statements even when they refer to the same tissue.

### Sequence and feature components

Chemical structures, therapeutic protein and antibody sequences and small-molecule physicochemical properties are represented by learned component nodes rather than by their raw strings or values (Table 4). SMILES strings of small molecules and of ADC linkers and payloads are neutralized and canonicalized with RDKit [53], without stereochemistry, and embedded with the ChemBERTa-2 chemical language model [54]. Therapeutic protein chains are embedded with the ESM-2 protein language model [55], and each antibody heavy chain is joined to its paired light chain by a GGGGS linker before embedding, with an alternate pair of a bispecific antibody treated as a separate input. For physicochemical features, up to 37 DrugBank properties, including molecular mass, lipophilicity, polar surface area, hydrogen-bond counts, ionization and solubility, are retained when observed for at least 75% of molecules with nonzero variance, log-transformed when strongly right-skewed, standardized and imputed by distance-weighted k-nearest neighbors. Language-model embeddings are mean-pooled over tokens. The vectors of each family are min– max scaled on the training set and factorized by non-negative matrix factorization (NMF) [56] as implemented in scikit-learn [57]. An input is linked to every factor whose loading reaches a factor-specific threshold, the (1 − p) quantile of positive training loadings, or to its highest-loading factor when none qualifies, so a compound can carry several components of one family. Raw strings, values and embeddings are stored separately from the graph. A component node therefore encodes membership in a learned factor: interventions linked to the same component received the same factor assignment, which does not imply a shared chemical substructure, sequence motif or physical subunit, and stereoisomers share the same SMILES input.

**Table 4.** Component families. Embeddings or normalized feature vectors are min–max scaled and factorized by non-negative matrix factorization (NMF). An input is assigned to every factor whose loading reaches the (1 − p) quantile of that factor’s positive training loadings, or to its highest-loading factor when none qualifies; factors with fewer than seven positive training loadings have no threshold. Encoders are ChemBERTa-2 [54] and ESM-2 [55].

| Component family | Input | Encoder | NMF factors | Top fraction p | Relation from Compound |
| --- | --- | --- | --- | --- | --- |
| SMILES sequence | Canonical SMILES of small molecules and of ADC linkers and payloads | ChemBERTa-2 (77M, masked language model) | 40 | 0.10 | Has SMILES Sequence Component; Linker or Payload SMILES Sequence Component (ADC) |
| Protein sequence | Therapeutic protein chain sequences | ESM-2 (650M) | 20 | 0.10 | Protein Sequence Component |
| Antibody sequence | Heavy chain joined to its paired light chain by a GGGGS linker | ESM-2 (650M) | 40 | 0.10 | Antibody Sequence Component |
| Small-molecule features | Normalized DrugBank physicochemical properties | None (numeric vector) | 8 | 0.15 | Has Features Component |

### Relation mapping

The graph distinguishes targeting, binding, inhibition, activation, modulation, metabolism and transport relationships between compounds and their molecular targets. These relation families preserve differences between molecular recognition, functional action and processes influencing drug disposition. In particular, association with a metabolic enzyme or transporter carries a different pharmacological meaning from inhibition of a therapeutic target.

Source relation labels are mapped to the predicates registered for each source–target node-type pair in the relation registry. A semantic relation mapper, which queries a large language model (OpenAI GPT-4.1) rather than embeddings, selects among the predicates permitted for the node-type pair and can return several predicates or none. Its cached output forms a predicate-mapping table of 601 rules (Supplementary Data 1), which was reviewed manually but not systematically validated. The table specifies the predicate or predicates, if any, for each source relation label and node-type pair (Fig. 4c): 426 rules map to a single predicate; 90 decompose the label into several predicates, so that an agonist assertion yields both Binds and Activates and a blocker yields Binds and Inhibits; and 85 are not mapped, namely 69 ontology or cross-reference relations without an equivalent in the schema, 11 unspecified or ambiguous compound actions or uses, and 5 gene-level labels, four of them negated GO annotations. The same mapper aligns relations proposed during enrichment. Each transformed edge keeps its source origin, such as DrugBank or GWAS Catalog, and a reference to the extracted triplet from which it was derived. Duplicate observations are merged into a single edge whose source origins and triplet references are collected in a list, which feeds the source-support signal of confidence scoring, whereas conflicting action labels from different sources are retained without adjudication. Differences between source assertions may reflect assay conditions, biological context, terminology or conflicting findings; their interpretation therefore depends on both the relation type and the supporting evidence.

### Review-gated enrichment of investigational therapeutics

Investigational therapeutics are often sparsely represented in structured resources. An on-demand, AI-assisted enrichment workflow, implemented in an internal curation application, therefore proposes information about a focal drug and relevant comparators (Fig. 2). Web-search-enabled language models (OpenAI gpt-4o-search-preview and gpt-5-search-api, both used over the course of this work) search for relation-specific information, including approved, investigational and off-label indications, contraindications, molecular targets and their actions, metabolizing enzymes, transporters, adverse events, pathways, GO concepts, cell types, anatomical sites and half-life, together with modality-specific properties such as SMILES strings, antibody format and chain sequences, protein subunit sequences and the antibody, linker and payload of ADCs. Field-specific instructions request canonical terms: MeSH terms for diseases, Uberon terms for anatomy, GO-style terms for GO concepts, Cell Ontology terms for cells, HGNC symbols for genes and MedDRA preferred terms for adverse events. For each proposed relation, the prompt requires a verbatim quotation of at most about 25 words that names both the drug and the related entity, with omissions marked, together with its source URL; responses are parsed and deduplicated by entity and relation label, and quotations are not automatically verified against the source page. Proposed entities and predicates are aligned to the graph vocabularies and relation schema with the same node and relation mappers used in construction; off-label indications are mapped to Indicated for, because the schema has no separate off-label predicate. The workflow is tuned for recall rather than precision, and searches are not restricted to a fixed evidence date. Prompt templates and the suggestion-file structure are provided in Supplementary Data 2.

Enrichment output is saved as a suggestion file, separate from the graph, so generating suggestions changes neither the graph nor any learned representation. Reviewers inspect the supporting quotations and links, accept or omit each suggestion, edit attributes or add concepts manually, and save a reviewed drug file; suggestions that could not be mapped to a standard identifier are not offered for acceptance. A reviewed file can be used in two ways: an embedding can be induced with a pretrained GNN encoder whose weights are held fixed, or the file can be merged into a project graph on which models are retrained. The reviewed information therefore defines what a representation-learning model knows about a new therapeutic. Because sparse neighborhoods may reflect limited study rather than erroneous biology, the evidence supporting each asserted relationship, rather than connectivity or co-occurrence, is central to review.

## Data Records

### Graph composition

The graph contains 166,261 nodes of 27 types and 8,106,906 directed edges of 129 relation types (Table 5). Node counts are distinct endpoint identifiers and edge counts are rows of the edge table, including the reciprocal rows added for Gene–Gene genetic associations. The node types comprise compounds; biomedical entities, namely genes, diseases, phenotypes, side effects, pathways, GO concepts, anatomical structures and cells; classification nodes, namely ATC classes, drug categories, Drug Types, chemical taxonomy and the vaccine, gene-therapy, cell-therapy and protein-therapy type taxonomies; modality-specific descriptor nodes, namely SMILES sequences, small-molecule feature components, antibody and protein sequences, antibody formats and genetics, heavy- and light-chain isotypes and linker types; and half-life bins. The edge table has seven columns, holding the identifier, name and type of the source and target nodes and the relation, which is serialized as predicate::source type:target type (for example, Has Half Life::Compound:Half Life). Modality-specific relations are stored with Compound as their source type, because the therapeutic categories are scopes of a single Compound identity.

**Table 5.** Composition of the graph. Counts are computed from the edge table of the post-processed base graph; compounds are a subset of all nodes. Percentages use the denominator of 18,429 compounds; categories are mutually exclusive because each compound has exactly one Drug Type, and Other comprises compounds whose therapeutic category could not be resolved during construction. A companion drug inventory lists 18,430 interventions; one vaccine without an edge in the post-processed graph is excluded.

| Quantity | Count | Definition or share |
| --- | --- | --- |
| Nodes | 166,261 | Distinct endpoint identifiers in the edge table |
| Node types | 27 | Node types observed in the edge table |
| Directed edges | 8,106,906 | Edge-table rows, including reciprocal Gene–Gene rows |
| Relation types | 129 | Relation types observed in the edge table |
| Source resources | 40 | Resources used for vocabularies or relation extraction (Table 3) |
| Compounds | 18,429 | Compound nodes, approved and experimental |
| *Compounds by therapeutic category* |  |  |
| Small molecules | 13,815 | 74.96% |
| Monoclonal antibodies | 1,235 | 6.70% |
| Protein-based therapies | 1,210 | 6.57% |
| Vaccines | 547 | 2.97% |
| Gene therapies | 304 | 1.65% |
| Antibody–drug conjugates | 253 | 1.37% |
| Cell therapies | 205 | 1.11% |
| Other | 860 | 4.67% |
| Total | 18,429 | 100.00% |

The 18,429 compounds span eight mutually exclusive therapeutic categories, because each compound has exactly one Drug Type (Table 5; Fig. 5a). Small molecules account for 74.96% of compounds, followed by monoclonal antibodies (6.70%) and protein-based therapies (6.57%); vaccines, gene therapies, ADCs and cell therapies extend the resource beyond small-molecule pharmacology. The categories describe coverage, whereas modality-specific relations describe molecular organization: small molecules, monoclonal antibodies, protein therapies and ADCs carry molecular-composition descriptors; vaccines, gene therapies and cell therapies carry a type classification only; and compounds in the Other category, whose therapeutic category could not be resolved during construction, have no modality-specific relations (Fig. 5b). This distinction separates the presence of a therapeutic class in the graph from the depth of its modality-specific characterization, which matters when comparing the abundant small molecules with less numerous biologic and composite therapies.

**Figure 5.**
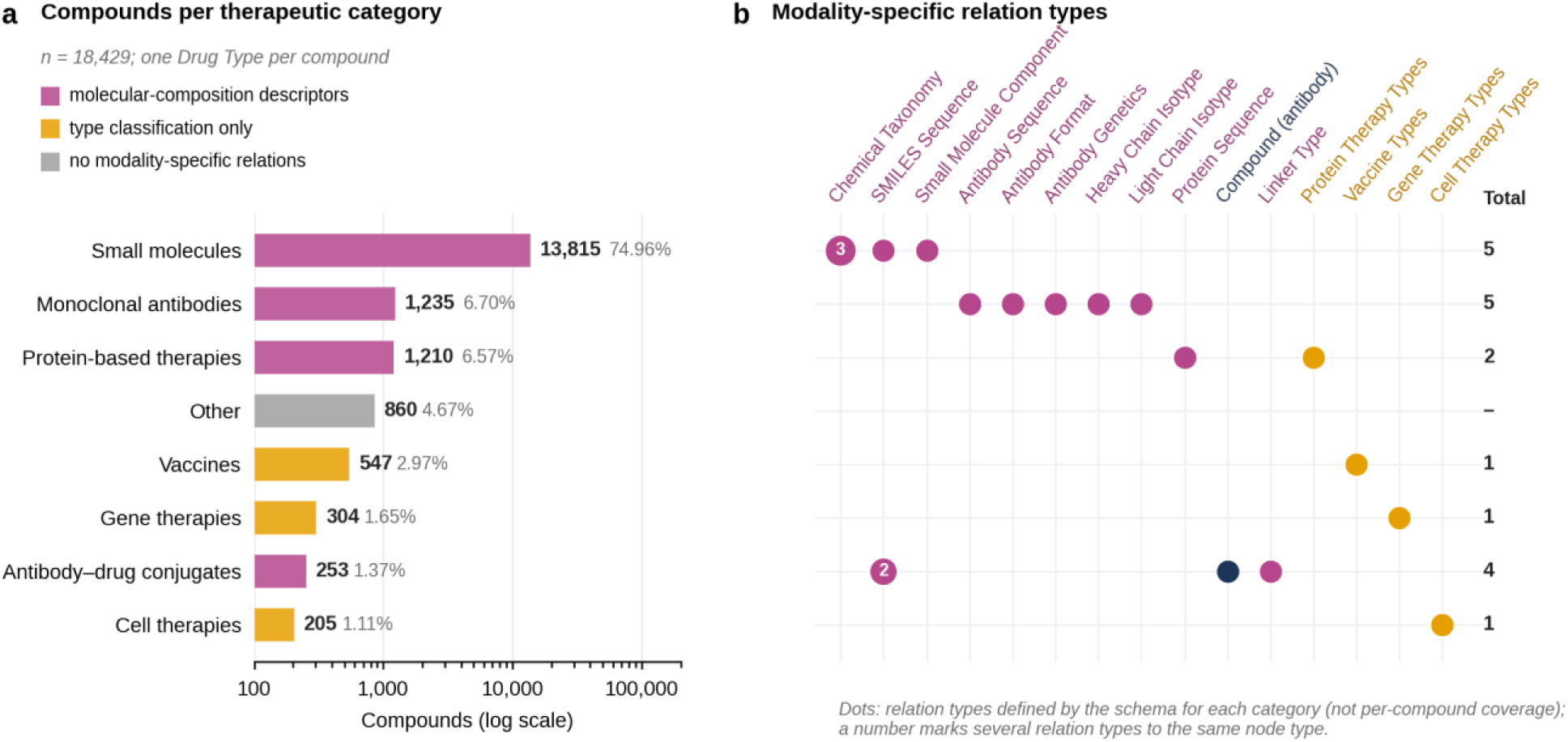
Therapeutic composition and modality-specific descriptors. **a**, Compounds per therapeutic category (log scale), coloured by the depth of modality-specific characterization defined by the schema. **b**, Modality-specific relation types defined by the schema for each category; numbers indicate several relation types to the same node type, such as class, subclass and superclass for chemical taxonomy or payload and linker components for ADC SMILES sequences. The matrix shows schema definitions rather than per-compound coverage.

### Relation schema

Table 2 lists the compound relation schema and the resources that populate it; all 48 compound relation types are observed in the graph. The remaining 81 relation types connect biological and clinical entities, for example through gene–gene interactions, gene–disease associations, gene expression in tissues and cells, and ontology hierarchies. Table 6 details the eight direct compound-to-context relations connecting drugs to pathways, GO concepts, anatomy and cells. Each compound has exactly one Compound::Drug Type::Has Type relation, which assigns its therapeutic category and determines the applicable modality-specific relations. Compound–disease relations distinguish indications (Indicated for) from investigational uses (Investigated for), contraindications and adjunct, symptomatic or preventative use. Indicated for denotes a source-asserted indication, which in enrichment also covers off-label use, rather than a regulatory approval status, which requires separate evidence.

**Table 6.** Direct compound–context relations. The eight relations connect Compound nodes to pathways, GO concepts, anatomy and cells. Operational definitions follow the text-extraction instructions, which the enrichment workflow shares (Supplementary Data 2). All eight relations are text-derived from Wikipedia and PubMed; compound–pathway relations also draw on Reactome, where pathway participation is mapped to Acts Through.

| Relation identifier | Biological domain | Operational definition |
| --- | --- | --- |
| Compound::Pathway::Acts Through | Pathway | Pathway through which the compound exerts its primary mechanism |
| Compound::Pathway::Activates | Pathway | Pathway activated or stimulated by the compound |
| Compound::Pathway::Inhibits | Pathway | Pathway inhibited or blocked by the compound |
| Compound::GO::Enables | GO molecular function | Molecular function enabled or performed by the compound itself |
| Compound::GO::Involved In | GO biological process | Biological process in which the compound participates, excluding clinical outcomes and disease states |
| Compound::GO::Located In | GO cellular component | Cellular component in which the compound's molecular function takes place |
| Compound::Anatomy::Targets | Anatomy | Anatomical structure in which the compound exerts its primary pharmacological action, excluding sites of metabolism or excretion |
| Compound::Cell::Targets | Cell | Cell type directly targeted by drug action |

### Compound identity and shared pharmacology

Shared compound relationships include molecular targets, indications, contraindications, adverse effects, drug categories and ATC classes at four levels. These connections provide a common neighborhood for interventions with different physical architectures. Molecular targets are represented at the gene level, with NCBI Gene as the standard vocabulary. Target-action relations distinguish binding from activation or inhibition, while metabolic and transporter associations connect the therapeutic to processes affecting its disposition.

The distinction between target identity and target action is central to drug comparison. Two compounds associated with the same target may act in different directions, and a target association may reflect metabolism or transport rather than the intended therapeutic mechanism. Shared pharmacological classes provide a further level of comparison, while disease and adverse-effect relationships connect molecular information to clinical context.

### Modality-specific descriptions

Small-molecule representations link each compound to a chemical taxonomy derived from DrugBank and to two families of learned components, SMILES sequence components and physicochemical feature components (Table 4). Chemical class, subclass and superclass relate molecules through structural categories, whereas shared components connect molecules whose encoded structures or property profiles received the same factor assignments: erlotinib and gefitinib, for example, share their taxonomy assignments, two SMILES sequence components and a feature component, as well as the EGFR target (Fig. 6). Components summarize encoded inputs rather than chemical fragments, and they provide inputs for models that combine chemical similarity with a drug’s biomedical neighborhood.

**Figure 6.**
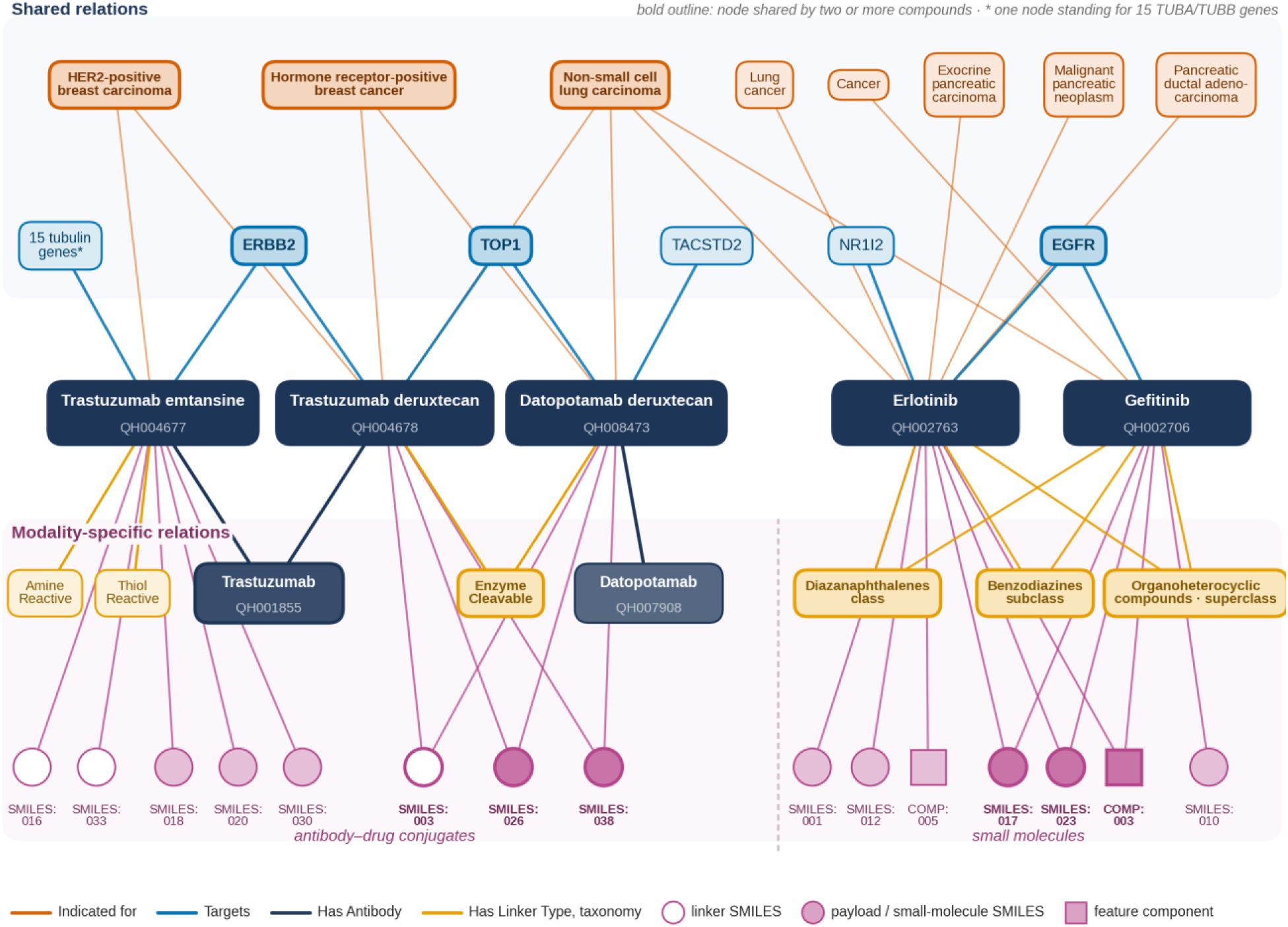
Example compound records. Subgraph of five compounds and their shared and modality-specific relations, taken from the graph. Antibody–drug conjugates (left): trastuzumab emtansine and trastuzumab deruxtecan share the trastuzumab antibody and the ERBB2 target but differ in linker type, linker and payload components and payload targets; trastuzumab deruxtecan and datopotamab deruxtecan share linker and payload components, the Enzyme Cleavable linker type and the TOP1 target but differ in antibody and antigen target. Small molecules (right): erlotinib and gefitinib share their chemical taxonomy, two SMILES sequence components, a feature component and the EGFR target. Non-small cell lung carcinoma links two ADCs and both small molecules. Intervention identifiers appear below compound names; SMILES:nnn and COMP:nnn are node labels, and identical labels denote the same node. Shared component nodes indicate shared assignments to learned factors (Table 4) rather than shared substructures. Indicated for denotes a source-asserted indication rather than regulatory approval status. *The tubulin node stands for the 15 tubulin genes targeted by trastuzumab emtansine (TUBA1A, TUBA1B, TUBA1C, TUBA3C, TUBA3E, TUBA4A, TUBB, TUBB1, TUBB2A, TUBB2B, TUBB3, TUBB4A, TUBB4B, TUBB6 and TUBB8).

Monoclonal-antibody representations link each antibody to antibody sequence components, learned from its paired heavy and light chains (Table 4), and to its format, genetics descriptors, heavy-chain isotype and light-chain isotype, drawn from SAbDab and ADCdb. Origin-related categories, such as human or humanized, and format-related descriptions characterize the therapeutic construct alongside its target relationships, so that antibodies with related binding targets retain distinct molecular descriptions within the shared graph. Protein-based therapies are represented through protein sequence components learned from their chain sequences and a protein-therapy type, drawn mainly from DrugBank, which characterize the therapeutic beyond a generic target label.

ADCs are represented as composite interventions with antibody, linker and payload information drawn from ADCdb. Has Antibody links the ADC to the Compound node of its monoclonal antibody, which carries its own antibody descriptors; payload SMILES components, assigned by the same SMILES model as small molecules, describe the chemical cargo; and linker SMILES components and linker type describe the connection between them. The intact Compound retains the intervention’s biological and clinical associations, including targets contributed by both antibody and payload, such as ERBB2 and TOP1 for trastuzumab deruxtecan. In the graph, trastuzumab deruxtecan and trastuzumab emtansine share the trastuzumab antibody and the ERBB2 target but differ in linker type, linker and payload components and payload targets (TOP1 versus 15 tubulin genes), whereas trastuzumab deruxtecan and datopotamab deruxtecan share their linker and payload components and the TOP1 target but differ in antibody and antigen target (ERBB2 versus TACSTD2; Fig. 6). Similarity in one component therefore does not imply equivalence of the complete drug’s activity or exposure. Vaccines, gene therapies and cell therapies are currently characterized by type classifications from internally defined taxonomies, without molecular-composition descriptors.

### Pharmacokinetic and pharmacodynamic context

Half-life annotations add an exposure-related dimension to the drug descriptions. Half-life values from DrugBank are converted to hours and summarized for each compound as the median of terminal or elimination half-life mentions, or of all numeric mentions when none is labeled terminal or elimination. Each summary value is assigned to one of 12 internally defined bins with inclusive lower and exclusive upper bounds: less than 30 minutes, 30–60 minutes, 1–2, 2–4, 4–8, 8–12 and 12–24 hours, and 1–3, 3–7, 7–14, 14–30 and 30–45 days. The last bin is labeled ‘>30 days’ in the graph, and values of 45 days or more receive no half-life node. Eleven ordering relations link consecutive bins from shorter to longer duration, so comparisons between nonadjacent bins follow a path along this chain. Half-life annotations cover 2,151 of the 18,429 compounds (11.67%), from 15.54% of protein-based therapies and 12.97% of small molecules to less than 1% of cell therapies, vaccines and compounds in the Other category (Table 7); ten small molecules carry two bins. These annotations can distinguish drugs with related target biology but different persistence in the body. They do not specify the kinetic phase, active metabolite, population or dose regimen of an individual value, and they complement molecular structure and qualitative action relations rather than specifying a concentration–time or exposure–response model. For cell therapies and biologics without a conventional elimination half-life, the enrichment workflow requests the time to half of maximal expansion instead, so enriched half-life annotations are not uniformly elimination half-lives.

**Table 7.** Half-life coverage by therapeutic category. A compound is covered when it has at least one Has Half Life edge. Ten small molecules have two half-life bins, so edges exceed covered compounds by ten. Percentages use the compounds of each category as the denominator.

| Therapeutic category | Compounds | With half-life | Coverage | Half-life edges |
| --- | --- | --- | --- | --- |
| Small molecules | 13,815 | 1,792 | 12.97% | 1,802 |
| Monoclonal antibodies | 1,235 | 133 | 10.77% | 133 |
| Protein-based therapies | 1,210 | 188 | 15.54% | 188 |
| Vaccines | 547 | 2 | 0.37% | 2 |
| Gene therapies | 304 | 16 | 5.26% | 16 |
| Antibody–drug conjugates | 253 | 13 | 5.14% | 13 |
| Cell therapies | 205 | 2 | 0.98% | 2 |
| Other | 860 | 5 | 0.58% | 5 |
| Total | 18,429 | 2,151 | 11.67% | 2,161 |

Binding, inhibition, activation and related action labels describe qualitative pharmacology. Associations with pathways, cells and anatomy extend this description beyond the immediate target to the context in which drug action occurs. Edge rows hold only endpoint identifiers, names and types and the relation, without dose, population, assay conditions or potency, so the graph does not provide quantitative PK/PD measurements; quantitative modeling would additionally require measurements such as exposure, affinity, potency or response, interpreted under defined experimental and physiological conditions [11].

### Multiscale biological and clinical context

Direct drug–pathway, drug–GO, drug–cell and drug–anatomy relationships make biological context accessible in a compound’s local neighborhood. They complement paths that reach the same concepts through molecular targets: interventions acting on different targets may still converge on a common pathway or cellular process, providing a basis for mechanistic comparison beyond target overlap.

GO supplies biological-process, molecular-function and cellular-component concepts [22]. Drug–GO relationships use these concepts to describe pharmacological associations, whereas conventional GO annotations characterize gene products. In the graph, drug–GO, drug–cell and drug–anatomy relations are text-derived, and the extraction instructions define them by the compound’s own activity: for example, a molecular function performed by the compound rather than by its target protein, or the anatomical site of its primary pharmacological action rather than a site of metabolism or excretion (Table 6). The interpretation of a drug–GO edge therefore depends on its relation semantics and supporting evidence, and a graph path connects relevant observations without, by itself, establishing a causal mechanism.

Disease, symptom, phenotype and adverse-effect entities connect therapeutic descriptions to clinical outcomes and liabilities. Anatomy and cell concepts provide physiological context, while pharmacological classifications group interventions by shared characteristics. The same information can support hypothesis generation and predictive modeling, although the role of each relation depends on the scientific task.

## Technical Validation

### Structural validation

Quality control operates mainly on the transformed, harmonized triplets rather than on raw source data. Structural checks verify that each extract satisfies its contract, namely required fields, recognized source and target node types and approved identifier vocabularies, and that each transformed relation uses a predicate registered for its node-type pair. These checks establish the structure of the extracted data, not the biological correctness of individual assertions.

### Confidence scoring of compound–gene and compound–disease assertions

Automated confidence scoring is applied to transformed triplets from Compound to Gene or Disease, excluding contraindications (Fig. 7a). Four signals on a [0, 1] scale contribute to a weighted score: the number and provider diversity of source-origin records, through a bounded transform (weight 0.30); local graph connectivity, measured by Adamic–Adar similarity [58] through a separate bounded transform (0.20); PubMed co-occurrence, scored as 1 − (1 − max(NPMI, 0))³, where NPMI is the normalized pointwise mutual information [59] computed from Entrez query counts [33] (0.20); and a language-model assessment of biological plausibility (0.30). A scored triplet is marked valid when its weighted score exceeds 0.35 or its plausibility score exceeds 0.45 (Fig. 7b), so plausibility enters both through its weight and through a separate threshold, and a sufficiently high plausibility score is decisive on its own. Triplets outside the scored relation families remain eligible for inclusion in drug records without this score-based check. Because PubMed counts were obtained from queries and caches rather than a frozen literature corpus, and because co-occurrence and plausibility do not verify the stated relationship, these scores serve as screening signals rather than independent verification of an assertion.

**Figure 7.**
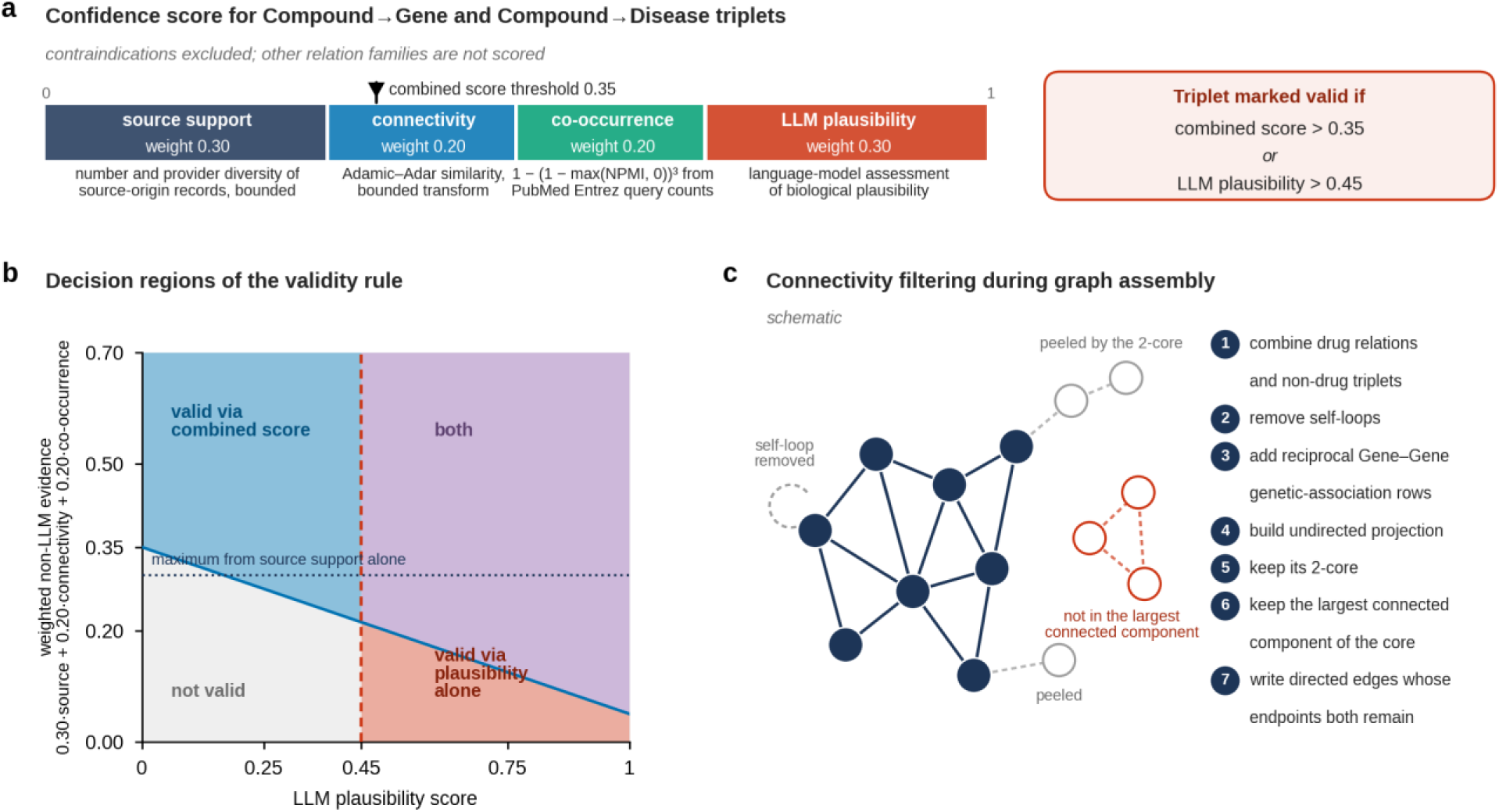
Confidence scoring and connectivity filtering. **a**, Signals and weights of the confidence score applied to Compound→Gene and Compound→Disease triplets (contraindications excluded), and the validity rule. **b**, Decision regions of the validity rule, plotted against the language-model plausibility score and the weighted contribution of the other three signals, each bounded to [0, 1]. Triplets with a plausibility score above 0.45 are valid regardless of the other signals; the dotted line marks the largest contribution that source support can make alone. **c**, Schematic of connectivity filtering during assembly: self-loops are removed, nodes outside the 2-core of the undirected projection are peeled away, and only the largest connected component of the core is retained.

### Connectivity filtering during graph assembly

Graph assembly applies structural filters (Fig. 7c). Drug records are scored for coverage across relation categories before assembly, and the loader combines the selected drug relations with non-drug triplets, removes self-loops and adds reciprocal rows for Gene–Gene genetic associations. It then constructs an undirected projection, retains nodes in its 2-core [60], selects the largest connected component within that core and writes the directed edges whose endpoints remain in that component. Because the 2-core retains only nodes with at least two neighbors within the core, sparsely annotated compounds and entities can be absent from the base graph; such interventions can instead be represented through enrichment.

### Known issues in transformation

Two properties of the transformation step affect the graph described here and are disclosed rather than corrected. First, five of the 601 predicate-mapping rules assign Gene Ontology annotations that carry a NOT qualifier to positive Gene–GO predicates (Enables, Involved in or Located in), so the Gene–GO edges derived from them state the opposite of the source annotation. Second, conflicting action labels from different sources, such as Inhibits and Activates for the same compound–gene pair, are retained without adjudication. Because each edge keeps a reference to the extracted triplet from which it was derived, edges affected by either issue can be identified and filtered; the affected rules are flagged in Supplementary Data 1.

### Review of identities and enrichment suggestions

Compound identity assignments were checked by manual, qualitative review, and enrichment suggestions enter project graphs only after expert review of their supporting quotations and sources. Together with the automated checks above, these procedures screen assertions and graph structure but do not estimate error rates. Text-derived relations, including all direct compound–context relations, are subject only to structural checks and assembly filters, and an edge absent from the graph should not be read as a verified negative.

## Usage Notes

Relations should be read through their semantics and provenance. Indicated for denotes a source-asserted indication rather than regulatory approval, and enrichment maps off-label uses to the same predicate. Compound–GO, compound–cell and compound–anatomy relations describe the compound’s own activity as defined by the extraction instructions (Table 6) rather than gene-product annotations, and they are text-derived and unscored. Because the anatomy and cell instructions also accept statements about receptor expression and affected cell populations, these relations mark sites associated with drug action rather than verified sites of action. Reactome-derived Acts Through edges represent pathway participation rather than a primary mechanism. Half-life annotations are categorical, and enriched values for cell therapies and some biologics denote time to half-maximal expansion rather than elimination half-life. Half-life bins cover values below 45 days, and the bin labeled ‘>30 days’ spans 30 to under 45 days.

Absence of an edge or descriptor denotes missing information rather than a biological negative. Descriptor coverage differs by therapeutic category (Fig. 5b), and the 2-core filter removes sparsely connected entities, so similarity between interventions can reflect annotation density as well as biology. When indications or other compound relations are prediction targets, related features such as ATC classes, drug categories and text-derived context can encode the target information and should be controlled for. Gene–GO edges derived from negated annotations and compound–gene pairs that carry conflicting action labels should be filtered or reviewed in analyses that depend on them (Technical Validation). Shared sequence or feature components indicate similar encoded inputs rather than shared substructures or motifs (Table 4).

Enrichment suggestion files are unreviewed proposals and are not graph content. A reviewed drug file can be used with a pretrained encoder whose weights are held fixed, which computes embeddings for the enriched drugs only, or merged into a project graph on which models are retrained, which can change all learned representations. Analyses that compare drugs across these settings should record which path was used.

## Discussion

We have introduced a drug-centric knowledge graph, which represents the therapeutic intervention as a structured object embedded in a shared biomedical network. A common Compound identity connects diverse therapies through pharmacological, biological and clinical relationships, while modality-specific descriptions preserve molecular and compositional differences. Half-life annotations add an exposure-related dimension, and direct pathway, GO, cell and anatomy relations extend the immediate context beyond molecular targets.

The contribution lies in coordinating these dimensions within a drug-centric representation. Prior graphs already provide rich drug features, mechanistic paths and pharmacological information; the design described here organizes shared therapeutic identity and specialized molecular descriptions together. This is particularly relevant to biologics and composite therapies, for which a target label or a single chemical structure gives an incomplete account of the intervention, and it supports comparisons at both whole-drug and component levels while retaining common biological context (Fig. 6). The three principles, namely intervention-level identity, a two-tier schema of shared and modality-specific relations, and relation semantics that separate recognition, action and disposition, are not tied to particular sources or implementations and could be adopted by other biomedical graphs, including those assembled with general-purpose frameworks such as BioCypher [9].

The organization also has implications for representation learning. Descriptor nodes change graph connectivity, while sequences, chemical structures and pharmacological attributes can add information beyond connectivity. Distinguishing these contributions requires controlled comparisons that separate topology from feature content and treat modality-specific, biological-context and pharmacokinetic information as distinct factors, rather than assuming that greater graph detail necessarily improves a model.

Several limitations define the scope of the resource. Molecular descriptions vary across therapeutic classes, and a class’s presence in the graph does not imply uniform descriptor coverage. Sources were not selected through a documented protocol, inclusion filters are source-specific rather than governed by a common quality policy, and text-derived relations, including the direct compound–context relations, were not subject to confidence cutoffs or automated scoring. Language models contribute to vocabulary and predicate mapping, plausibility scoring and enrichment; their outputs have not been quantitatively validated, the predicate-mapping table received only manual review, and the validity rule allows a plausibility score alone to admit an assertion. Five mapping rules invert negated GO annotations, and conflicting action labels are retained without adjudication. Compound-identity rules were assessed only by qualitative review. The 2-core filter favors well-connected entities and can exclude sparsely annotated interventions from the base graph. The PK description emphasizes half-life annotations and does not constitute a quantitative PK/PD model. Component models and half-life binning are described from the construction code, and the fitted component models of the reported build were not re-inspected; text extraction also asks the language model to check consistency with a compound’s known indication, which can introduce knowledge beyond the source text. Execution records for the reported build were not retained, so the build cannot be reproduced exactly, and the graph is proprietary, which limits independent access. An assertion-level audit estimating accuracy by modality, relation family, source and enrichment status would address the most important of these gaps.

Future development can extend the graph with context-qualified pharmacological observations, in which a numerical value such as clearance, exposure or potency is linked to its parameter, units, biological system and dosing or assay conditions, as illustrated by PK-DB [61]. Source manifests recorded at acquisition, frozen literature snapshots for co-occurrence scoring and retained execution records would strengthen reproducibility, and publicly shareable schema definitions and example subgraphs would support comparison with other resources. The drug-centric design provides a concrete basis for studying whether jointly representing what a therapeutic is, how it acts and where it acts improves drug representations for biomedical research.

## Data availability

The graph is a proprietary resource of QuantHealth and is not publicly distributed with this article. The compound relation schema, source inventory and predicate-mapping rules are provided as Supplementary Data 1.

## Code availability

The graph was constructed with custom Python notebooks and package functions executed on Databricks; this code and the drug-embedding software are proprietary and not publicly available. Job-level execution records for the reported build were not retained, so its exact software environment cannot be reconstructed. The text-extraction and enrichment prompt templates and the suggestion-file structure are provided as Supplementary Data 2.

## Author contributions

Study concept and design: Alon Bar, Michael Shapiro, Amichai Perlman, Flavio Dormont, Omri Matalon.

Methodology: Alon Bar, Gilad Erez, Doron Levinson, Tal Raisfeld, Uri Shamgar, Michael Shapiro.

Data curation and analysis: Alon Bar, Gilad Erez, Doron Levinson, Tal Raisfeld, Flavio Dormont.

Software development: Alon Bar, Uri Shamgar, Gilad Erez, Doron Levinson, Tal Raisfeld.

Validation: Alon Bar, Gilad Erez, Michael Shapiro.

Preparation of figures and visualizations: Flavio Dormont, Alon Bar.

Drafting of the manuscript: Flavio Dormont, Amichai Perlman, Alon Bar, Michael Shapiro, Omri Matalon.

Project administration: Michael Shapiro, Omri Matalon.

Supervision: Michael Shapiro, Flavio Dormont, Omri Matalon.

## Ethics statement

This article describes a biomedical knowledge resource and does not report an analysis of individual-level patient data.

## Supplementary information

**Supplementary Data 1.** Compound relation schema, source inventory and predicate-mapping rules.

**Supplementary Data 2.** Text-extraction and enrichment prompt templates, field-specific instructions and suggestion-file structure.

