## Supplementary Data 2 for "Drug-centric knowledge graph: representing therapeutic modality, pharmacology and multiscale biology"

### Supplementary Data 2. Text-extraction and enrichment prompt templates, field-specific instructions and suggestion-file structure

*Supplement to: Drug-centric knowledge graph: representing therapeutic modality, pharmacology and multiscale biology.*

#### Overview

This document reproduces the prompt templates used to extract compound–context relations from text during base-graph construction (Section 1) and the prompt template and field-specific instructions of the enrichment workflow (Sections 2–5), together with the structure of the suggestion file that the enrichment workflow produces (Section 6). It records configured prompts rather than a verified record of any particular extraction or enrichment run, and it does not show that extracted or suggested facts are accurate.

Prompt text, casing, spelling and punctuation are reproduced verbatim, including typographical errors in the original, except for the antibody-property instructions in Section 4, which are given as intended (see the note there). Text in braces, such as {compound_name} or {drug_name}, is inserted at run time.

Enrichment relation searches insert the drug name, a short drug-type summary as context and a field-specific instruction (Section 3) into the general relation-extraction prompt (Section 2), which is sent to a web-search-enabled model (OpenAI gpt-4o-search-preview or gpt-5-search-api; both have been used over the course of this work). Responses are parsed into items with an entity, a relation label, a supporting quotation and a URL, and are deduplicated by entity and relation label; parsing does not verify the quotation against the source page. Drug-type and property searches (Section 4) use separate search procedures, each of which adds its own output-format and evidence instructions.

#### 1. Base-graph text extraction

Compound–pathway, compound–GO, compound–anatomy and compound–cell relations of the base graph are extracted with OpenAI GPT-4.1 at temperature 0, using structured output. For each available context, three messages are sent: the shared instruction below as a developer message, the context as a user message and one task instruction as a second user message. A context is either the English Wikipedia article for the compound, when one is found, with its title, URL, summary and full text, or a batch of up to five PubMed abstracts with their identifier, title, journal and year; up to ten abstracts are retrieved per compound by relevance-ranked search without a publication-date restriction. The three task groups are processed separately for each context, and their outputs are merged. These are the retrieval settings of the construction code rather than a record of the articles processed.

##### *Shared developer instruction*

```
You are a medical knowledge extraction system specializing in drug mechanism of action (MOA)
analysis.

Your role is to extract structured relationships between drugs and biological entities from
scientific text sources including Wikipedia and PubMed abstracts.
```

```
Key principles:
- Extract ONLY information explicitly stated in the provided text
- Focus on the specific drug mentioned in the extraction task
- Use standard ontology terminology (Gene Ontology, Uberon, Cell Ontology, Reactome pathways)
when possible
- Distinguish between direct drug effects vs. background information
- Provide text references to support your extractions
- Return structured JSON output as specified

You will receive:
1. Source text (Wikipedia or PubMed abstracts)
2. Specific extraction task instructions
```

##### *Pathway task*

```
=== EXTRACTION TASK ===

DRUG: {compound_name}

OBJECTIVE:
Extract ALL explicitly stated OR directly inferred pathway relationships involving the drug
{compound_name}.
Prioritize completeness while maintaining strict textual grounding.

TASK:
Identify and extract every pathway that the text explicitly states is:
- activated by
- inhibited by
- used as a mechanistic route by
the drug {compound_name}.

--------------------------------
RELATION TYPES
--------------------------------

1. activates
Definition:
Pathways whose activity or signaling is increased, stimulated, or promoted by {compound_name}.

Trigger phrases include (not exhaustive):
"activates", "stimulates", "enhances", "upregulates",
"induces", "promotes", "augments", "leads to activation of",
"results in activation of"

2. inhibits
Definition:
Pathways whose activity or signaling is reduced, blocked, or suppressed by {compound_name}.

Trigger phrases include (not exhaustive):
"inhibits", "blocks", "suppresses", "downregulates",
"attenuates", "prevents activation of", "negatively regulates",
"reduces signaling through"

3. acts_through
Definition:
Pathways explicitly described as mediating, enabling, or constituting the mechanism of action
of {compound_name}.

Trigger phrases include (not exhaustive):
"acts through", "mediated by", "via", "through",
"mechanism involves", "dependent on", "requires activation of",
"effects are mediated by"

EXTRACTION PRINCIPLES:

1. Compound grounding
- Extract ONLY relationships where {compound_name} is the causal agent.
- Ignore pathways associated with other drugs, ligands, or general biology.

2. Explicit textual evidence
```

```
- The relationship must be directly stated in the text.
- Do NOT infer pathway involvement from downstream genes, proteins, or phenotypes unless the
pathway itself is explicitly named.

3. Context window
- The pathway must appear:
  a) in the same sentence as {compound_name}, OR
  b) in an immediately adjacent sentence where {compound_name} is clearly the grammatical or
referential subject (including pronouns or abbreviations).

4. Coreference handling
- Treat abbreviations, synonyms, or references (e.g., "the drug", "this compound") as
referring to {compound_name} if unambiguous in context.

5. Exclusions
IGNORE pathways mentioned only in:
- Database annotation lists (GO, KEGG, Reactome summaries, etc.)
- Cross-references or supplementary tables
- Background or review-style descriptions not explicitly tied to {compound_name}

6. Pathway validity
- Pathways must represent recognized biological pathways:
  signaling, metabolic, immune, cellular stress, cell-cycle, or transcriptional pathways.
- Use standardized pathway names whenever possible.

OUTPUT CONTRACT:
Return a JSON object with "extractions" array. Each extraction has:
{
  "extractions": [
    {
      "relation": "activates" | "inhibits" | "acts_through",
      "values": [string, ...],
      "text_reference": string | null
    },
    ...
  ]
}

RULES:
- Group pathways by their relation type
- Use concise pathway names
- If no pathways found for a relation type, omit that extraction
- If nothing found at all, return {"extractions": []}
```

##### *Gene Ontology task*

```
=== EXTRACTION TASK ===

DRUG: {compound_name}

TASK: Extract ALL GO relationships for the drug {compound_name} across multiple relation types
and GO categories.

RELATION TYPES:
1. enables -  Molecular Function that {compound_name} enables or performs
    Molecular Functions represent molecular-level activities performed by gene products, such
as “catalysis” or “transcription regulator activity”.
    Molecular Functions correspond to activities that can be performed by individual gene
products (i.e. a protein or RNA), but some activities are performed by molecular complexes
composed of multiple gene products, when the activity cannot be ascribed to a single gene
product of the complex.
    Examples of broad functional terms are "catalytic activity" and "transporter activity".
    Examples of more specific functional terms are "adenylate cyclase activity" or "insulin
receptor activity".
   - Trigger phrases: "enables", "has molecular function", "exhibits activity", "catalyzes",
"binds to"

2. involved_in - Biological Process that {compound_name} is involved in or participates in
    Biological Process are the larger processes or programs accomplished by the concerted
action of multiple molecular activities.
    Examples of broad BP terms are DNA repair or signal transduction.
```

```
    Examples of more specific terms are cytosine biosynthetic process or D-glucose
transmembrane transport.
   - Trigger phrases: "involved in", "participates in", "role in process", "affects process",
"regulates"
   - Do not include clinical outcomes or disease states (e.g., "cancer progression",
"reduction of heart failure symptoms")

3. located_in -  Cellular Component where the molecular function of {compound_name} takes
place
    Cellular Component serves to capture the cellular location where the molecular function of
{compound_name} the takes place.
    CCs include:
     - cellular anatomical structures, encompassing cellular entities such as the plasma
membrane and the cytoskeleton, as well as membrane-enclosed cellular compartments such as the
mitochondrion.
     - stable protein-containing complexes of which they are parts.
   - Examples: "mitochondrion", "cell membrane", "nucleus"
   - Trigger phrases: "located in", "localizes to", "found in", "accumulates in",
"compartment", "site of action"
   - refrain from extracting organ/tissue locations where the drug is absorbed, metabolized,
or excreted (e.g., "intramuscular injection site", "liver", "kidney")

EXTRACTION PRINCIPLES:
- Include only information about the requested compound {compound_name}, ignore GO terms
related to other compounds or general background
- ONLY extract GO terms where the text explicitly describes the compound's DIRECT biological
activity
- GO terms must be in the SAME SENTENCE or IMMEDIATE CONTEXT as the compound name and its
action
- IGNORE GO terms mentioned in:
  * Database annotation lists (e.g., "GO annotations include:", "RGD annotations:", "UniProt
annotations:")
  * Cross-reference sections without mechanistic context
  * Lists of terms without connecting verbs or actions
- GO terms should describe what the COMPOUND does, not what its target protein does
- Verify the GO term is RELEVANT to the compound's known therapeutic or biological class
- If a GO term seems unrelated to the compound's primary function, exclude it

FIELD INSTRUCTIONS:
- Use GO-style terminology when possible
- Prioritize specificity over generality (e.g., prefer "adenylate cyclase activity" over
"catalytic activity")
- Prioritize GO terms that appear in sentences with action verbs connecting them to the
compound

OUTPUT CONTRACT:
Return a JSON object with "extractions" array. Each extraction has:
{
  "extractions": [
    {
      "relation": "enables" | "involved_in" | "located_in",
      "values": [string, ...],
      "text_reference": string | null
    },
    ...
  ]
}

RULES:
- Group GO terms by their relation type
- Use GO-style terminology
- Keep Molecular Functions, Biological Processes, and Cellular Components separate
- If no GO terms found for a relation type, omit that extraction
- If nothing found at all, return {"extractions": []}
```

##### *Anatomy and cell target task*

```
=== EXTRACTION TASK ===

DRUG: {compound_name}
```

```
TASK: Extract ALL target relationships for the drug {compound_name}, including anatomical and
cellular targets.

RELATION TYPES:
1. **target_anatomy** - Anatomical structures or organ systems where {compound_name} exerts
its PRIMARY PHARMACOLOGICAL ACTION
   - Use Uberon ontology terminology (e.g., "brain", "bone marrow", "gastrointestinal tract",
"mucosa of stomach", "sheath of Schwann")
   - Focus on the site of action where the drug produces its therapeutic effect
   - Trigger phrases: "site of action", "therapeutic target", "acts on", "exerts effects in",
"receptor expression in"

2. **target_cell** - Specific cell types that {compound_name} TARGETS
   - Use Cell Ontology (CL) terminology (e.g., "hepatocyte", "cardiomyocyte", "neoplastic
cell", "pancreatic ductal cell", "polymodal nociceptor")
   - Include both therapeutic target cell types and affected cell populations
   - Trigger phrases: "targets", "selective for", "acts on", "cell-type specific", "expressed
in", "affects"

EXTRACTION PRINCIPLES:
- Include only information about the requested compound {compound_name}, ignore targets
related to other compounds or general background
- ONLY extract targets where the text explicitly describes the compound's ACTION on that
anatomy/cell
- Targets must be in the SAME SENTENCE or IMMEDIATE CONTEXT as the compound name and its
pharmacological action
- IGNORE targets mentioned in:
  * Database annotation lists
  * Pharmacokinetic/ADME contexts (metabolism, clearance, elimination)
  * Cross-reference sections without mechanistic description
- For anatomical targets: Distinguish between sites of therapeutic action vs. sites of
metabolism/excretion
- For cell targets: Extract cells that are the intended therapeutic target, not incidental
affected cells
- Verify the target is CONSISTENT with the compound's known therapeutic indication

FIELD INSTRUCTIONS:
- For anatomical targets: Focus on pharmacological action sites, not metabolic/clearance sites
- For cell targets: Prefer using the most specific definition available in the text
- Use appropriate ontology terminology (Uberon for anatomy, CL for cells)
- Prioritize targets that appear with action verbs describing the compound's effect

OUTPUT CONTRACT:
Return a JSON object with "extractions" array. Each extraction has:
{
  "extractions": [
    {
      "relation": "target_anatomy" | "target_cell",
      "values": [string, ...],
      "text_reference": string | null
    },
    ...
  ]
}

RULES:
- Keep anatomical and cellular targets separate by using correct relation types
- Use ontology-style terminology
- If no targets found for a relation type, omit that extraction
- If nothing found at all, return {"extractions": []}
```

#### 2. Enrichment: general relation-extraction prompt

```
You are given a drug and some context.

Drug: {drug_name}
Context: {context}
Instruction: {instruction}
```

```
Task:
1) Find entities related to the drug ("relations") according to the instructions.

2) For each relation, assign a concrete relation_type label that is justified by evidence
   in a reliable source (e.g., indicated_for, inhibits, binds, targets, involved_in_pathway,
   off_label_use, investigational_indication, activates, metabolized, causes_side_effect
etc.).

3) Provide a verbatim quote (exact words + skipping "..." if needed) from the source URL that:
   (a) clearly supports BOTH the *relation* and the chosen *relation_type*.
   (b) skips parts of the quote irrelevant to the *relation* and the *relation_type* (using
"...").
   (c) **does not include any other entity**. If the source sentence lists other entities,
remove them and their context, but **insert ("...") in their place** to clearly mark the
omission, while preserving grammar and enough context to support the relation_type.
   (d) The quote must be **at most two short sentences or ~25 words**, using ("...")
aggressively to remove unnecessary text.
   (e) **Do not elide the drug name or the relation text**. Both must appear verbatim in the
quote if present in the source.


   **Examples:**
   Acceptable:
   - relation: "CCR7"
   - Source_text: "It also shows activity in β-arrestin assays against CCR6, and some activity
at CCR7 and CXCR2."
   - reasoning: "It also shows ... some activity at CCR7 ..."

   Unacceptable:
   - "It also shows activity in β-arrestin assays against CCR6, and some activity at CCR7 and
CXCR2." (mentions multiple entities)
   - "It has activity at CCR7." (paraphrase, not verbatim)
   - Any quote longer than two sentences or containing unnecessary context.

4) Include the URL where the quote was taken.

Formatting:
Return ONLY a JSON object (no extra text, no markdown fences) with this schema:

{
  "items": [
    {
      "relation": "entity string (without the drug name)",
      "relation_type": "LABEL",
      "reasoning": "\"exact verbatim quote from the page supporting the LABEL\"",
      "url": "https://example.com/source"
    }
  ]
}

Rules:
- Use trustworthy sources when possible (e.g., labels, guidelines, reputable references).
- When text is removed, always insert ("...") where the removal occurs, to indicate omitted
content.
- Each item must have all four fields and the quote must directly justify the relation_type.
- Prefer concise, highly relevant quotes (one or two sentences).
- The "reasoning" must contain the drug's name and the relation string (no elision of either).
- Do not place ("...") at the start or end of a quote — use it only within sentences.
```

#### 3. Enrichment: field-specific relation instructions

Each instruction below is inserted into the general prompt as {instruction}. For GO and pathway searches, a shared instruction block is appended to each field-specific instruction.

#### Disease

##### *Approved indications*

```
Identify at least 10 approved indications for the drug.
Search in clinical / regulatory reliable sources.
Use canonical MeSH terms at the most specific level available for the disease.
Do not include parent categories; expand them into all child diseases as separate relations.
```

##### *Investigational indications*

```
Identify at least 10 investigational disease indications for the drug.
Search through reliable sources for phase I-III studies.
Use canonical MeSH disease terms at the most specific level available.
Do not return parent categories; expand into children diseases, as separate relations - and
match each to the parent reasoning and url.
```

##### *Off-label indications*

```
Find all OFF-LABEL disease indications for the drug, supported by real-world or case-based
clinical uses outside regulatory approval, case reports, retrospective series, or
observational studies in credible sources.
Return each disease in canonical MeSH form.
```

##### *Contraindications*

```
Identify ALL official contraindications *for the given drug*.
Search FDA product labeling (Contraindications section).
Include both absolute and relative contraindications.
Return each contraindicated condition in canonical MeSH form.
Exclude warnings, precautions, or adverse effects that are not true contraindications.
Do not include exclusion criteria from clinical trials as contraindications.
```

#### Anatomy

##### *Anatomical targets*

```
Identify ALL known ANATOMICAL targets of the drug.

*target_anatomy* - Anatomical structures or organ systems where the drug exerts its PRIMARY
PHARMACOLOGICAL ACTION

Guidelines:
- Use Uberon ontology terminology (e.g., "brain", "bone marrow", "gastrointestinal tract",
"mucosa of stomach", "sheath of Schwann")
- Focus on the site of action where the drug produces its therapeutic effect
- Trigger phrases for example: "site of action", "therapeutic target", "acts on", "exerts
effects in", "receptor expression in"
- Distinguish between sites of therapeutic action vs. sites of metabolism/excretion
- Focus on pharmacological action sites, not metabolic/clearance sites
- for all anatomy relations you find, determine searcher_relation_type = targets

Scope rules:
- ONLY extract targets where the text explicitly describes the compound's ACTION on that
anatomy
- Include only direct anatomical targets of action.
- Do NOT include diseases, cell types, pathways, or proteins unless they clearly refer to an
anatomical structure.
- If the drug targets specific anatomical regions or substructures, include each separately.
```

```
Be inclusive rather than restrictive — if an anatomical structure is clearly described as
being directly targeted or affected, include it.
```

#### Gene Ontology

##### *Shared GO instructions (appended to each GO instruction)*

```
EXTRACTION PRINCIPLES:
- Include only information about the requested compound, ignore GO terms related to other
compounds or general background
- ONLY extract GO terms where the text explicitly describes the compound's DIRECT biological
activity
- IGNORE GO terms mentioned in:
  * Database annotation lists (e.g., "GO annotations include:", "RGD annotations:", "UniProt
annotations:")
  * Cross-reference sections without mechanistic context
  * Lists of terms without connecting verbs or actions
- GO terms should describe what the COMPOUND does, not what its target protein does
- Verify the GO term is RELEVANT to the compound's known therapeutic or biological class
- If a GO term seems unrelated to the compound's primary function, exclude it

FIELD INSTRUCTIONS:
- Use GO-style terminology when possible
- Prioritize specificity over generality (e.g., prefer "adenylate cyclase activity" over
"catalytic activity")
- Prioritize GO terms that appear in sentences with action verbs connecting them to the
compound
```

##### *Molecular functions (enables)*

```
Identify all the **molecular functions** that the compound enables or performs.
Molecular Functions represent molecular-level activities performed by gene products, such as
“catalysis” or “transcription regulator activity”.
Molecular Functions correspond to activities that can be performed by individual gene products
(i.e. a protein or RNA), but some activities are performed by molecular complexes composed of
multiple gene products, when the activity cannot be ascribed to a single gene product of the
complex.
Examples of broad functional terms are "catalytic activity" and "transporter activity".
Examples of more specific functional terms are "adenylate cyclase activity" or "insulin
receptor activity".
- Trigger phrases: "enables", "has molecular function", "exhibits activity", "catalyzes",
"binds to".

Output constraint for "relation":
- The "relation" MUST be a molecular-function activity phrase (an activity enabled/performed),
not just an entity name.
- It must be written as an activity noun phrase, typically "<entity> <activity>", where
<activity> is a function term such as "binding", "receptor binding", "catalytic activity",
"transporter activity", "enzyme activity", etc.
- Do NOT output bare entities (e.g., "X" is invalid). Always output an activity phrase (e.g.,
"X binding" is valid).

Mapping rule:
- searcher_relation_type must be 'enables' for all items

Quote constraint:
- The reasoning quote must explicitly support the activity phrasing (contain the drug name and
the relevant activity trigger, e.g., bind/inhibit/catalyze/activate + the entity).
```

##### *Biological processes (involved in)*

```
Identify all the biological process that the compound is involved in or participates in.
Biological Process are the larger processes or programs accomplished by the concerted action
of multiple molecular activities.
```

```
Examples of broad BP terms are DNA repair or signal transduction.
Examples of more specific terms are cytosine biosynthetic process or D-glucose transmembrane transport.
- Trigger phrases: "involved in", "participates in", "role in process", "affects process", "regulates"
- Do not include clinical outcomes or disease states (e.g., "cancer progression", "reduction of heart failure symptoms")

Mapping rule:
- searcher_relation_type must be 'involved_in' for all items
```

##### *Cellular components (located in)*

```
Identify all the cellular components (CC) where the molecular function of the compound takes place.
Cellular Component serves to capture the cellular location where the molecular function of the compound takes place.
CCs include:
  - cellular anatomical structures, encompassing cellular entities such as the plasma membrane and the cytoskeleton, as well as membrane-enclosed cellular compartments such as the mitochondrion.
  - stable protein-containing complexes of which they are parts.
- Examples: "mitochondrion", "cell membrane", "nucleus"
- Trigger phrases: "located in", "localizes to", "found in", "accumulates in", "compartment", "site of action"
- refrain from extracting organ/tissue locations where the drug is absorbed, metabolized, or excreted (e.g., "intramuscular injection site", "liver", "kidney")

Mapping rule:
- searcher_relation_type must be 'located_in' for all items
```

#### Pathway

##### *Shared pathway instructions (appended to each pathway instruction)*

```
EXTRACTION PRINCIPLES:
- Include only information about the requested compound, ignore pathways related to other compounds or general background
- ONLY extract pathways where the text explicitly describes the compound's ACTION on that pathway
- IGNORE pathways mentioned in:
  * Database annotation lists (e.g., "GO annotations include:", "RGD annotations:")
  * Cross-reference sections
  * General background information not specific to this compound
- Pathways should be directly stated as targets or path of the pharmacoligal activity of the drug, not inferred from related proteins or genes

FIELD INSTRUCTIONS:
- Use standard pathway nomenclature (e.g., "PI3K/AKT signaling", "NF-kappa-B pathway", "mTOR pathway")
- Include signaling pathways, metabolic pathways, cellular pathways
- Be specific about which relation type applies to each pathway
```

##### *Activates*

```
Extract ALL pathway activation relationships for the compound.
activates - Pathways that the compound ACTIVATES or STIMULATES
  - Trigger phrases: "activates", "stimulates", "upregulates", "enhances", "promotes pathway", "induces pathway"

Mapping rule:
- searcher_relation_type must be 'activates' for all items
```

##### *Inhibits*

```
Extract ALL pathways that the compound INHIBITS or BLOCKS
   - Trigger phrases: "inhibits", "blocks", "suppresses", "downregulates", "prevents pathway
activation"

Mapping rule:
- searcher_relation_type must be 'inhibits' for all items
```

##### *Acts through*

```
Extract ALL pathways through which the compound exerts its primary mechanism
   - Trigger phrases: "acts through", "mechanism involves", "mediated by pathway", "via
pathway", "through pathway"

Mapping rule:
- searcher_relation_type must be 'acts_through' for all items
```

#### Cell

##### *Cellular targets*

```
Identify ALL known CELL targets of the drug.

*cell targets* - Specific cell types that the drug TARGETS.
   - Use Cell Ontology (CL) terminology (e.g., "hepatocyte", "cardiomyocyte", "neoplastic
cell", "pancreatic ductal cell", "polymodal nociceptor")
   - Include both therapeutic target cell types and affected cell populations
   - Trigger phrases examples: "targets", "selective for", "acts on", "cell-type specific",
"expressed in", "affects"

Guidelines:
- Extract cells that are the intended therapeutic target, not incidental affected cells
- Prefer using the most specific definition available in the text
- Prefer established biological cell type names (e.g., T cell, B cell, epithelial cell,
neoplastic cell, macrophage).
- Include each distinct cell type separately.

Scope rules:
- Include only direct cellular targets of action.
- Do NOT include diseases, pathways, proteins alone, or tissues unless they clearly refer to a
cell type.
- If the drug has multiple cellular targets, list each separately.

Be inclusive rather than restrictive — if a cell type is clearly described as being directly
targeted or affected, include it.
```

#### Gene

##### *Direct targets*

```
Identify all known GENE or PROTEIN targets of the drug.
Focus on **direct molecular targets** that the drug binds to or modulates to produce its
pharmacologic effect.
Search authoritative pharmacology and bioinformatics sources.
For each target, return the **canonical gene symbol** (HGNC-approved symbol, e.g., EGFR,
DRD2).
If the drug has multiple subunit targets or complexes, include each gene separately.
```

##### *Binding*

```
Identify all GENES whose protein products are known or predicted to **bind** the drug
molecule, based on experimental or computational evidence.
Include both **primary pharmacologic targets** and **secondary/off-target bindings** supported
by binding affinity data (e.g., Kd, Ki, IC50, EC50) or structural/docking studies.
Search authoritative pharmacologic and bioactivity sources such as DrugBank, ChEMBL,
BindingDB, IUPHAR/BPS, PubChem BioAssay, UniProt, or peer-reviewed binding studies.
Return the **canonical HGNC gene symbol** for each bound protein.
Aim to include both high-affinity and well-documented off-target bindings when available.
```

##### *Inhibition, activation and modulation*

```
Identify all genes whose protein products are directly or indirectly **inhibited**,
**activated**, or **modulated/regulated** by the drug at the molecular level.
Focus on experimentally demonstrated pharmacologic actions, such as receptor or enzyme
inhibition, activation, or allosteric modulation.
Search authoritative mechanistic sources.

Define relation_types as:
- **inhibits** — the drug blocks or decreases the activity of the target (e.g., enzyme
inhibitor, receptor antagonist, ion-channel blocker).
- **activates** — the drug increases or triggers target activity (e.g., receptor agonist,
enzyme activator, channel opener).
- **modulates/regulates** — the drug alters target activity or expression without full
activation or inhibition (e.g., allosteric modulator, transcriptional regulator, partial
agonist/antagonist).

Return the **canonical HGNC gene symbol** for each target.
Exclude targets without mechanistic evidence or with only indirect genetic associations.

If multiple mechanisms are described for the same gene (e.g., partial agonist and modulator),
include each distinct mechanism as a separate relation with its own supporting quote and URL.
```

##### *Metabolizing enzymes*

```
Identify the genes whose encoded enzymes **metabolize the drug** (i.e., the drug is a
substrate). Include Phase I and Phase II enzymes.
Search authoritative pharmacokinetic sources.

Return the **canonical HGNC gene symbol** for each enzyme.
Prefer **specific isoforms** over families. If a source only names a family/subfamily (e.g.,
“CYP3A”), do **not** infer isoforms; include only isoforms explicitly supported by the quoted
source.

Scope:
- Include **major and clinically relevant minor** metabolizing enzymes of the **parent drug**.
If an **active metabolite** is formed and its formation enzyme is specified, include that
enzyme as well (still as drug → gene “metabolized_by”; the quote may mention the metabolite).
- For **prodrugs**, enzymes catalyzing **bioactivation** are in scope (still
“metabolized_by”).

Aim to capture **all major pathways** and **notable minor routes** with solid evidence.
```

##### *Transporters and carriers*

```
Identify the genes encoding **transporter proteins** that actively or passively **transport
the drug** across cellular membranes — i.e., the drug is a **substrate** of the transporter.
Search authoritative pharmacokinetic and transporter sources.
Return the **canonical HGNC gene symbol** for each transporter. Prefer **specific isoforms**
when available; if only a family (e.g., “OATP”) is mentioned, include only explicitly named
members.

Scope:
- Include **uptake and efflux transporters** that move the **parent drug** or **active
```

```
metabolites** across intestinal, hepatic, renal, or blood–brain barriers.
- Both **active transporters (ABC family)** and **facilitated transporters (SLC family)** are
in scope.
- Include transporters with **in vitro**, **in vivo**, or **clinical pharmacokinetic**
evidence that the drug is a substrate.

Aim to capture **all major uptake and efflux transporters** known to handle the drug,
prioritizing those with strong experimental or regulatory evidence.
```

##### Side effect

###### *Adverse events*

```
Identify the **clinical side effects (adverse events)** of the drug, as documented in
authoritative sources.
Sources to prioritize: **FDA/EMA product labeling** (Adverse Reactions; Warnings and
Precautions; Boxed Warning), clinical trial publications/meta-analyses, and high-quality AE
databases (e.g., DrugBank, SIDER, FAERS analyses).
Avoid "fda.gov/drugs/development-approval-process-drugs/drug-approvals-and-databases".

Return each side effect using a **standardized clinical term** (prefer **MedDRA Preferred
Term**; MeSH acceptable if MedDRA not available).

Labeling for relation_type:
- Use **BOXED_WARNING** for toxicities explicitly listed in the boxed warning.
- Otherwise use **ADVERSE_EVENT** for clinically recognized side effects.

Scope:
- Include **on-label** adverse reactions (from trials/label) and **serious risks** highlighted
in warnings.
- Laboratory abnormalities may be included **only if** recognized as adverse reactions in
labeling (e.g., “neutropenia”).
- If the source provides frequency or severity, prefer a quote that includes it, but keep
relation_type as above.

Exclude:
- **Contraindications**, **precautions** without an adverse event, and non-causal
associations.
- Symptoms of the **underlying disease** unless the drug increases their incidence.

Normalization:
- Use **canonical casing/spelling** for the AE term (e.g., “Interstitial lung disease”,
“Neutropenia”, “Nausea”).
- Avoid umbrella catch-alls when a **more specific MedDRA PT** is available (e.g., prefer
“Interstitial lung disease” over “Pneumonitis” only if the source uses ILD; otherwise match
the source term exactly).

Aim to capture **all major and clinically meaningful** adverse events, with emphasis on
**serious or common** reactions supported by explicit, verbatim evidence from the source.
```

#### 4. Enrichment: drug-type and property searches

##### Drug type

The drug-type search uses the evidence template below. The extracted type phrase is then mapped to the internally defined compound subtypes by a separate mapping prompt. {code_rule} asks for a company pipeline or research-and-development source when the drug name is a recognized investigational code name.

```
You are a biomedical expert.
```

```
        Task: For the drug "{drug_name}", return {target_count} high-quality public source(s)
that clearly describe what KIND of drug it is (its modality/type).

        Guidelines:
        - The snippet must explicitly name "{drug_name}" (or an official synonym) and indicate
its type (e.g., Small Molecule, Monoclonal Antibody, Antibody Drug Conjugate, CAR-T, gene
therapy, vaccine, etc.).
        - Prefer reliable sources: FDA, EMA, DrugBank, PubMed, ChEMBL, company site
(labels/inserts, press or pipeline pages). Avoid forums and generic scraped sites.
        {code_rule}
        {note_clause}
        - Return concise 1–3 sentence snippets.

        Output JSON (object with a single field "evidences" = array of exactly {target_count}
object(s)):
        {
        "evidences": [
            {
            "source_url": "<direct URL>",
            "snippet": "<1–3 sentence excerpt>",
            "extracted_phrase": "<short noun phrase like 'Small Molecule', 'Monoclonal
Antibody', 'Antibody Drug Conjugate', etc.>"
            }
            {comma_example}
        ]
        }
```

#### General and small-molecule properties

##### *Half-life*

```
What is the half-life time of the drug? Search in a clinical reliable source.If the drug is a
cell therapy or biologic where a conventional pharmacokinetic half-life is not defined,
instead return time to half of maximal expansion.
```

##### *Small-molecule SMILES*

```
What is the *exact* SMILES code of this drug? Search in a clinically reliable source, and make
sure you quote PERCISELY.
```

Small-molecule numeric features are searched with an empty instruction string, and the search procedure supplies a list of 13 feature names. Drug category is also searched with an empty instruction string, whereas ATC classes and chemical taxonomy are assigned by mapping procedures rather than by an instruction string.

#### Antibody–drug conjugates

##### *Linker identity*

```
What is the exact chemical linker used in the antibody–drug conjugate (ADC)?Search
authoritative primary sources only (FDA or EMA label/SmPC, or peer-reviewed literature).Return
the most chemically specific linker name or structure designation explicitly used in the
source,not a functional or class description.Do NOT return high-level linker classes or
mechanismsunless the source provides no more specific chemical identifier.If multiple payload
attachments exist, report the linker type used in the approved product; if the source truly
does not specify the linker, return 'N/A'. Return only: - Linker name (exact chemistry) -
Cleavable or non-cleavableDo not return DAR, payload, or antibody details—only the linker.
```

##### *Payload identity*

```
What is the payload used in the antibody–drug conjugate (ADC)? Search in an authoritative
clinical source (FDA/EMA label, SmPC, or peer-reviewed literature). Return the specific
cytotoxic payload name or class (e.g., 'deruxtecan', 'MMAE', 'DM1') and its type or mechanism
of action (e.g., 'topoisomerase I inhibitor', 'microtubule inhibitor'). If multiple payloads
are mentioned, report the primary one used in the approved product. If the source truly does
not specify the payload, return 'N/A'. Do not return DAR, linker, or antibody details—only the
payload.
```

##### *Antibody identity*

```
What is the antibody used in the antibody–drug conjugate (ADC)? Search in an authoritative
clinical source (FDA/EMA label, SmPC, or peer-reviewed literature). Return only the generic
antibody name (INN) (e.g., 'Panitumumab'). If the ADC uses a bispecific or engineered
antibody, return the official INN and indicate the modification succinctly (e.g.,
“bispecific”).If the source truly does not specify the antibody, return 'N/A'. Do not return
DAR, linker, or payload details—only the antibody.
```

##### *Linker sequence*

```
What is the *exact* sequence-defined (specific) linker code of this ADC linker? make sure you
quote PERCISELY.
```

##### *Linker SMILES*

```
What is the *exact* SMILES code of this ADC linker sequence? make sure you quote PERCISELY.
```

##### *Payload SMILES*

```
What is the *exact* SMILES code of this ADC payload? Search in a clinically reliable source,
and make sure you quote PERCISELY.
```

##### *Linker type*

```
Determine the type of the provided linker, and select exactly one option from the list
```

#### Monoclonal antibodies

Antibody format, antibody genetics, heavy-chain isotype and light-chain isotype are each classified with a dedicated instruction, shown below as intended. In an earlier configuration of the workflow the antibody-format instruction was passed for all four properties; because each search was restricted to the valid options for the property concerned and the instruction was generic, the resulting classifications were not affected. Heavy- and light-chain sequences are searched with a dedicated chain-sequence prompt conditioned on the antibody format returned by the first classification.

##### *Antibody format*

```
Determine the antibody format of the drug and select exactly one option from the list
```

##### *Antibody genetics*

```
Determine the antibody genetics of the monoclonal antibody and select exactly one option from
the list
```

##### *Heavy-chain isotype*

```
Determine the heavy chain isotype of the monoclonal antibody and select exactly one option
from the list
```

##### *Light-chain isotype*

```
Determine the light chain isotype of the monoclonal antibody and select exactly one option
from the list
```

#### Other modalities

For type classifications, the allowed options are supplied with the instruction. Protein subunit sequences are searched with a dedicated subunit prompt rather than with the type instruction.

##### *Protein-therapy type*

```
Determine the Protein Based Therapy Type of the drug and select exactly one option from the
list
```

##### *Cell-therapy type*

```
Determine the Cell Based Therapy Type of the drug and select exactly one option from the list
```

##### *Gene-therapy type*

```
Determine the Gene Therapy Type of the drug and select exactly one option from the list
```

##### *Vaccine type*

```
Determine the Vaccine Type of the drug and select exactly one option from the list
```

### 5. Alignment and review of enrichment suggestions

Suggested relations are mapped to schema predicates by the relation mapper, which asks a language model to choose among the predicates permitted for the source–target node-type pair in the relation registry, or to return none. Disease, gene and side-effect suggestions pass the retrieved quotation to the mapper, whereas anatomy, GO, pathway and cell suggestions pass the searcher's relation label. Off-label indications are mapped to the Indicated for predicate, because the schema has no separate off-label predicate.

Target entities are resolved with the node mapper for the corresponding node type, which tries exact title matches and then semantic candidates with a language-model referee. A term may map to zero, one or several standard identifiers.

The workflow writes one suggestion file per drug. The curation application displays the candidate quotations and URLs and lets reviewers choose or edit fields before saving a separate, reviewed drug file. The prompts contain no fixed evidence-date cutoff, and runtime model settings and reviewed outputs require separate run records.

#### 6. Structure of the enrichment suggestion file

Each suggestion file describes one drug and is a reviewable proposal, not an accepted drug file or a graph update. The file name records the drug and project; these are not fields inside the file, and the file itself does not carry an intervention identifier.

The top-level fields are name (the requested drug name), brief (the search context), base_relations (shared Compound relations), drug_type (the selected modality and its specific suggestions) and count (a recursively computed entry count). base_relations is organized as target node type → predicate → list of proposed target nodes and can contain Drug Type, ATC, Drug Category, Disease, Anatomy, GO, Pathway, Cell, Gene, Side Effect and Half Life entries. A typical target entry contains id, name, type, origin, reasoning and url; fields vary by search, mappings can leave id empty, and some entries add confidence or based_on. Entries without an identifier are not offered for acceptance during review.

The drug_type branches depend on modality. Small molecules can have chemical-taxonomy, SMILES and numerical-feature suggestions; monoclonal antibodies can have format, genetics, isotype and chain-sequence suggestions; ADCs can have antibody and linker-type suggestions together with linker and payload SMILES; protein therapies can have subunit-sequence and type suggestions; and cell therapies, gene therapies and vaccines can have type suggestions. For sequences, raw_value holds the proposed sequence, and multi-chain suggestions can be lists with labels and pairing metadata. count counts selected nested entries; it is not a count of distinct nodes, verified facts or graph edges.

The following illustrative structure uses placeholders, not a real drug or a validated assertion.

```
{
  "name": "Example drug",
  "brief": "Short drug context",
  "base_relations": {
    "Drug Type": {
      "Has Type": [
        {"id": "TYPE_ID", "name": "Small Molecule", "type": "Drug Type"}
      ]
    },
    "Disease": {
      "Indicated for": [
        {
          "id": "DISEASE_ID",
          "name": "Disease name",
          "type": "Disease",
          "origin": ["ai_enrichment"],
          "reasoning": "Supporting explanation or quote",
          "url": "https://source.example"
        }
      ]
```

```
    }
  },
  "drug_type": {
    "id": "TYPE_ID",
    "name": "Small Molecule",
    "subtype_relations": {
      "Chemical Taxonomy": {
        "Class": [
          {"id": "CLASS_ID", "name": "Class name", "confidence": 0.8}
        ]
      }
    },
    "subtype_features": {
      "sequences": {
        "SMILES Sequence": {
          "raw_value": "SMILES_PLACEHOLDER",
          "value": null,
          "origin": ["ai_enrichment"],
          "method": null,
          "reasoning": "Supporting explanation",
          "url": "https://source.example"
        }
      },
      "Has SMILES Sequence Component": [
        {
          "raw_value": "SMILES_PLACEHOLDER",
          "value": null,
          "origin": ["ai_enrichment"],
          "method": null,
          "reasoning": "Supporting explanation",
          "url": "https://source.example"
        }
      ]
    }
  },
  "count": 4
}
```
